# *LINC01432* binds to CELF2 in newly diagnosed multiple myeloma promoting short progression-free survival to standard therapy

**DOI:** 10.1101/2024.06.27.600975

**Authors:** Richa Mishra, Prasanth Thunuguntla, Alani Perkin, Dhanusha Duraiyan, Katelyn Bagwill, Savannah Gonzales, Vanessa Brizuela, Steve Daly, Yoon Jae Chang, Mahdote Abebe, Yash Rajana, Kelly Wichmann, Catheryn Bolick, Jaiyana King, Mark Fiala, Julie Fortier, Reyka Jayasinghe, Mark Schroeder, Li Ding, Ravi Vij, Jessica Silva-Fisher

## Abstract

Multiple Myeloma (MM) is a highly prevalent and incurable form of cancer that arises from malignant plasma cells, with over 35,000 new cases diagnosed annually in the United States. While there are a growing number of approved therapies, MM remains incurable and nearly all patients will relapse and exhaust all available treatment options. Mechanisms for disease progression are unclear and in particular, little is known regarding the role of long non-coding RNAs (lncRNA) in mediating disease progression and response to treatment. In this study, we used transcriptome sequencing to compare newly diagnosed MM patients who had short progression-free survival (PFS) to standard first-line treatment (PFS < 24 months) to patients who had prolonged PFS (PFS > 24 months). We identified 157 differentially upregulated lncRNAs with short PFS and focused our efforts on characterizing the most upregulated lncRNA, *LINC01432*. We investigated *LINC01432* overexpression and CRISPR/Cas9 knockdown in MM cell lines to show that *LINC01432* overexpression significantly increases cell viability and reduces apoptosis, while knockdown significantly reduces viability and increases apoptosis, supporting the clinical relevance of this lncRNA. Next, we used individual-nucleotide resolution cross-linking immunoprecipitation with RT-qPCR to show that *LINC01432* directly interacts with the RNA binding protein, CELF2. Lastly, we showed that *LINC01432*-targeted locked nucleic acid antisense oligonucleotides reduce viability and increases apoptosis. In summary, this fundamental study identified lncRNAs associated with short PFS to standard NDMM treatment and further characterized *LINC01432,* which inhibits apoptosis.

***Key points:*** lncRNA expression was found to be dysregulated in patients with short PFS to standard multiple myeloma therapy. *LINC01432*-bound CELF2 inhibits apoptosis.

## Introduction

Multiple myeloma (MM) is a prevalent disease and is the fifteenth leading cause of cancer-related deaths in the United States.^1^ Despite the increasing availability of treatment regimens, nearly all patients with MM become refractory and die from the disease or its sequale.^2–4^ In addition, while some improvements in patient outcomes have been achieved using novel immunomodulatory agents, new approaches are still needed due to high toxicity and the development of drug resistance. ^5–7^ This underscores the need for new therapeutic approaches. Thus, knowledge surrounding the mechanisms and biomarkers of treatment resistance in MM patients are critically needed to support the development of novel MM therapies.

Long non-coding RNA (lncRNA) is defined as RNA greater than 200 nucleotides in length that is not translated into functional proteins. Prior studies have reported that lncRNAs can promote the pathogenesis of all cancer types,^8–11^ including MM.^12–18^ Many lncRNAs have also been shown to promote MM drug resistance, including *NEAT1, ANRIL, MEG3, LINC00461, H19*, and *PCAT1*.^12,19–28^ lncRNAs are expressed in the cytoplasm,^29^ the nucleus,^30,31^ and in other organelles, such as exosomes,^32,33^ and may be expressed in more than one subcellular location.^31^ The subcellular localization of a lncRNA is highly important and specific to its biological functions in the cell, which may include transcriptional regulation, translational regulation, and interaction with RNA binding proteins.^34^ As lncRNA expression is highly tissue specific, they hold promise as novel therapeutic targets that can be used as prognostic and diagnostic biomarkers.^35–38^ Further, recent advances in understanding the functions and crucial roles lncRNAs play in promoting cancer, including MM, increases their potential as targets for RNA-based therapeutics.^39–42^

Limited availability of RNA sequencing or single cell sequencing data from newly diagnosed multiple myeloma (NDMM) patients, along with the heterogeneity of treatment approaches used with MM patients, have hindered research on the global expression of lncRNAs in MM and characterization of their biological functions in response to current standard MM therapies. In this study, we used RNA sequencing data from a cohort of NDMM patients to identify lncRNAs that were associated with a short progression-free survival (PFS). We identified several lncRNAs that were highly upregulated in patients with short PFS, as compared to prolonged PFS, and determined that *LINC01432* bound to the RNA-binding protein, CELF2, to inhibit apoptosis and increase viability.

## Methods

### RNA sequencing data, patient samples, and cell lines

RNA sequencing data from NDMM patients were obtained from the Multiple Myeloma Research Foundation (MMRF) Clinical Outcomes in Multiple Myeloma to Personal Assessment of Genetic Profiles (CoMMpass) study (https://registry.opendata.aws/mmrf-commpass), accessed in February of 2021 (**Supplementary** Figure 1**, Supplementary Tables 1 and 2**). MM cell lines were generously provided by Dr. John DiPersio at Washington University in St. Louis (RPMI 8226, U266B1, MM1.S, and OPM2) and were all cultured in RPMI 1640 media (Invitrogen, Carlsbad, CA) supplemented with 15% fetal bovine serum (Invitrogen) and 1% penicillin/streptomycin (Invitrogen). MM1.R cell lines were purchased from ATCC (catalog number CRL-2975) and cultured in the same manner as the other cell lines. NDMM patient bone marrow aspirates were obtained from the Multiple Myeloma Tissue Banking Protocol (IRB 201102270) processed by the Siteman Cancer Center Tissue Procurement Core.

Full length *LINC01432* transcript was amplified via PCR and cloned into the pCFG5-IEGZ-GFP vector (generously provided by Dr. Chris Maher, Washington University in St. Louis, Piscataway, NJ) to create the pCFG5-IEGZ-GFP-Luc-*LINC01432* vector (pCFG5-*LINC01432*). Full vector length was confirmed by GeneScript. Retroviral infection of HEK 293T cells was performed by transfecting cells with 2µg of empty vector control or pCFG5-*LINC01432.* Transduction was conducted by harvesting viral supernatants and adding to U266B1 cells in the presence of 8µg/ml polybrene (Sigma), then centrifuged at 500g for three hours. Fresh media was then added and cells were sorted for positive GFP expression via flow cytometry. Cells containing virus expressing *LINC01432* or empty vector were selected for using 100µg/ml Zeocin. Validated cell lines showing high levels of *LINC01432* expression by RT-qPCR, as compared to empty vector, were used for subsequent assays.

*LINC01432* knockdown CRISPR/Cas9 cells were generated using the RPMI 8226 cell line. The sgRNAs were generated by the Genome Engineering and Stem Cell Center, Washington University in St. Louis. sgRNAs were cloned into the pLV hUbC-dCas9 KRAB-T2A-GFP plasmid (Addgene #672620). HEK 293T cells were infected with this lentivirus to induce expression of dCas9-KRAB,^43^ followed by transduction, similar as above into RPMI 8226 cells and validated knockdown of *LINC01432* expression via RT-qPCR.

### Transfection of locked nucleic acid antisense oligonucleotides

Locked nucleic acid GapmeR antisense oligonucleotides (LNA ASOs) targeting *LINC01432* (Qiagen, cat# 3653410) and CELF2 (Qiagen, cat# 339511), and negative control LNA ASOs (Qiagen, cat#148759394), were designed using the Qiagen Antisense LNA GapmeR Custom Builder (https://www.qiagen.com), sequences are listed in **Supplementary Table 3**. MM cells were seeded at a density of 500,000 cells/well in 6-well plates, transfected with respective ASOs at 50nM-100nM concentration using Lipofectamine 2000, and incubated for 48–72 hours. Cells were harvested and target knockdown was validated via RT-qPCR.

### RNA Sequencing data analyses

RNA sequencing data was processed and analyzed from the MMRF ComPASS study. Briefly, the GRCh37 reference genome (hs37d5 version)^44^ was used for assembly, then the data was integrated with supplementary contigs, including the complete repeating unit of ribosomal DNA, cancer-related viruses, and data from the external RNA Control Consortium. For gene and transcript models, Ensemble version 74 was used, along with additional annotations provided in the GTF file (https://github.com/tgen/MMRF_CoMMpass.git). For sequence alignment,STAR 2.3.1z (01/24/2013) ^45^ was used and BAM files were subsequently generated using SAMtools v.1.19. To ensure the rigor of these data, we performed quality control checks on the RNA BAM files using Picard RNA metrics and BamTools Ig Counts. DAVID^46^ was used to determine Gene Ontology and pathway analyses. POSTAR3^47^ was used to determine *LINC01432* RNA:protein binding.

### Multiplexed Fluorescent RNA in situ Hybridization (mFISH)

RNAScope was performed as previously described,^48^ with some modifications using RNAscope 2.5 HD Reagent Kit Red assay combined with Immunohistochemistry (Advanced Cell Diagnostics [ACD], Catalog #323180 and #322372) according to manufacturer’s instructions. Briefly, bone marrow aspirates or isolated tumors were applied to slides, baked in a dry air oven for one hour at 60°C, deparaffinized (Xylene for five minutes twice, followed by 100% ethanol for two minutes twice), hydrogen peroxide was applied for 10 minutes at room temperature, and co-detection target retrieval was performed using Steamer (BELLA) for twenty minutes and PBS-T washing. Slides were then incubated overnight with CUGBP2 (CELF2) antibody (Protein Tech, Cat#12921-1-AP) in a HybEz Slide Rack with damp humidifying paper and incubated overnight at 4°C. The next day, slides were washed in PBS-T then post-primary fixation was performed by submerging slides in 10% NBF for 30 minutes at room temperature. Slides were then washed with PBS-T and Protease Plus was added to each slide for 30 minutes at 40°C, then slides were washed with distilled water. Probes for *LINC01432* (ACD Cat# 878271) were then warmed at 40°C and hybridized with specific oligonucleotide probes for 2 hours at 40°C in HybEZ Humidifying System. RNA was then serial amplified and stained with Fast Red solution. Slides were blocked with co-detection blocker (ACD) for 15 minutes at 40°C and washed with PBS-T. Secondary Alexa Fluor 488 antibody (Abcam, cat#ab150081) was applied for one hour at room temperature in the dark. Finally, slides were washed with PBS-T, counter stained with DAPI (Sigma, cat#D9542) for 30 seconds, and mounted with ProLong Gold Antifade Reagent (Invitrogen, cat#P36930). Slides were imaged on the EVOS M5000 Imaging System (Invitrogen).

Analysis of mFISH and IHC images was performed by comparing expression of *LINC01432* or CELF2 between different cell lines or tissues and simultaneously verifying their cellular localization or intensity of expression. We first visualized our target RNA molecules using an EVOS M5000 imaging system and quantified targets with QuPath Software v0.5.1 to obtain cell count per region and number of spots per cell data. We then applied multiplex analysis followed by the cell distribution analysis for detecting lncRNA spots in each cellular compartment.

### In vitro phenotypic assays

We used the ApoTox-Glo Triplex Assay (Promega, Madison, WI) to simultaneously measure viability, cytotoxicity, and apoptosis in the same sample. We seeded 20,000 cells/well of Control CRISPR/Cas9, *LINC01432* knockdown CRISPR/Cas9, empty vector, *LINC01432* overexpression, or wild-type RPMI 8226 cells in triplicate into a 96-well plate at 100uls complete media per well. We began by first adding 20ul of Viability/Cytoxicity reagent to all wells, and briefly mixing for ∼30 seconds. Plates were then placed in a 37°C incubator for one hour. Next, we measured the intensity of fluorescence (relative fluorescence units) using 400Ex/505Em (viability) and 485Ex/520Em (Cytotoxicity) in Varioskan LUX microplate reader. To measure apoptosis, we next add 100ul Caspase-Glo 3/7 reagent to the same wells and briefly mixed by orbital shaking ∼20 seconds, followed by incubation at room temp for 30 minutes. Luminescence (relative luminescence units) was then measured using Varioskan LUX microplate reader to detect caspase activation.

We measured apoptosis by isolating cells and assessing via flow cytometry using BD Horizon V450 AnnexinV (BD Biosciences, Franklin Lakes, NJ). We seeded 500,000 cells/well in a 6-well plate for 24 hours. Cells were then harvested and the usual protocol was followed, per manufactures instructions. Briefly, cells were washed twice with PBS, then incubated V450 AnnexinV and Propidium Iodide (ThermoFisher) for fifteen minutes at room temperature in the dark. Apoptosis and DNA content was assessed on a flow cytometer machine (Novios, Becton Dickinson) by the Flow Cytometry Core of Siteman Cancer Center, Washington University in St. Louis. We collected a minimum of 50,000 cells per sample in triplicate. FlowJo Version 10 (Becton Dickinson) was used to analyze data.

### In vivo individual-nuhcleotide resolution cross-linking immunoprecipitation (iCLIP)

The iCLIP assay was performed as previously described.^31^ Briefly, cells were seeded at a density of twenty million cells/150 mm dish. The next day, cells were washed with cold PBS and media volumes were adjusted to 10ml/dish. Dishes were then uncovered and irradiated with 150 mJ/cm^2^ of UVA (254 nm) in a crosslinker device (Stratalinker). Cells were then harvested and centrifuged at 2000 RPM at 4°C for 5 minutes. Cell pellets were resuspended in 1ml of NP-40 lysis buffer (20mM Tris–HCl at pH 7.5, 100mM KCl, 5mM MgCl2, and 0.5% NP- 40) with 1µl protease inhibitor and 1mM DTT, incubated on ice for ten minutes, and then centrifuged at 10,000 RPM for 15 minutes at 4°C. Supernatants were collected, 1U/μl RNase T1 was added, then cell lysates were incubated at 22°C for 30 minutes. Protein G Beads were resuspended in 100µls NT2 buffer (50mM Tris–HCl at pH 7.5, 150mM NaCl,1mM MgCl2, 0.05% NP-40) with 5µg of respective antibodies, then rotated for one hour at room temperature. All antibodies are listed in **Supplementary Table 4**. Cell lysates were added to the beads and incubated for three hours at 4°C, the beads were washed with NT2 buffer, and then incubated with 20 units RNAse-free DNase I for 15 minutes at 37°C in a thermomixer, shaking slowly. Protein kinase buffer (141µls NP- 40 lysis buffer, 0.1% SDS, 0.5 mg/ml Proteinase K) was then added and incubated for 15 minutes at 55 °C in a thermomixer, shaking at maximum speed. Supernatants were then collected and RNA isolation was performed using a standard phenol:cholorform:isoamyl alcohol protocol. RNA was then reverse transcribed using SuperScript III First strand cDNA system, as per manufacturer’s protocol (ThermoFisher) and primers tiling *LINC01432* (**Supplemental Table 3**) were used to detect *LINC01432*:protein binding.

### In vivo myeloma models

All animal experiment protocols in this study were reviewed and approved by the Institutional Animal Care and Use Committee of Washington University in St. Louis. For subcutaneous injections, 2e^5^–1e^7^ cells (RPMI 8226 wild-type, U266B1 wild-type, Control CRISPR/Cas9, *LINC01432* knockdown CRISPR/Cas9, empty vector, or *LINC01432* overexpression) were subcutaneously injected NOD/SCID/γc^−/−^ (NSG) mice (N = 5–10 per group). Resulting tumor size was quantified weekly via caliper measurements, comparing length x width x height x 0.5. For post-analyses, subcutaneous tumor tissues were removed after sacrifice, formalin fixed, and paraffin embedded. This experiment was repeated twice.

### Data sharing statement

All RNA sequencing data is available at GEO under accession number GSE267013. All other data supporting the findings of this study are available within this article and its Supplementary Information files from the corresponding author, upon reasonable request.

## Results

### Identification of dysregulated lncRNAs with short progression-free survival to standard MM therapy

In order to identify lncRNAs that are differentially expressed in patients that exhibit a short progression-free survival (PFS) to the standard MM treatment approach, we analyzed transcriptome sequencing data from CD138+ bone marrow samples obtained from 115 NDMM patients in the MMRF CoMMpass study (**Supplementary** Figure 1). We assigned samples to one of two groups based on each patient’s length of PFS to standard MM therapy, short, those who had progression-free survival < 24 months (short PFS) from the first dose of MM treatment (N = 38), and prolonged progression-free survival, those with > 24 months PFS (N = 77), **Table 1, Supplemental Table 1**. We identified 157 upregulated and 91 downregulated lncRNAs in short PFS, as compared to prolonged PFS (log2Fold Change > +/-2, *p* < 0.05), **Figure 1a**. lncRNAs identified as being most differentially expressed in short PFS included *LINC01432, lnc-LGALS9B-7, LINC01916, Lnc-SPIDR-1, and MAGEA4-AS1*, **Figure 1b**. We also identified two lncRNAs previously reported to be associated with MM, MEG3^24,49,50^ , and H19,^51–54^ **Supplemental Table 1.** Next, we performed pathway analysis on all differentially expressed RNAs in short PFS to identify highly enriched gene sets associated with, but not limited to, staphylococcus aureus infection (*p* = 6.75e^-10^), transcriptional dysregulation in cancer (*p* = 1.05e^-06^), cytokine-cytokine receptor interactions (*p* = 1.42e^-05^), IL-17 signaling pathway (*p =* 2.31e^-05^), and ECM-receptor interactions (*p =* 5.5e^-05^), **Figure 1c**. Gene ontology analysis further showed high enrichment of more than one pathway associated with immune response, B cell mediated immunity, and multiple hemoglobin complexes, **Supplementary** Figure 2**, a and b**. This analysis of sequencing data from NDMM patients in the MMRF CoMMPass study allowed us to identify lncRNAs that are differentially expressed in patients that exhibit a short PFS to standard MM treatment.

**Figure 1.**
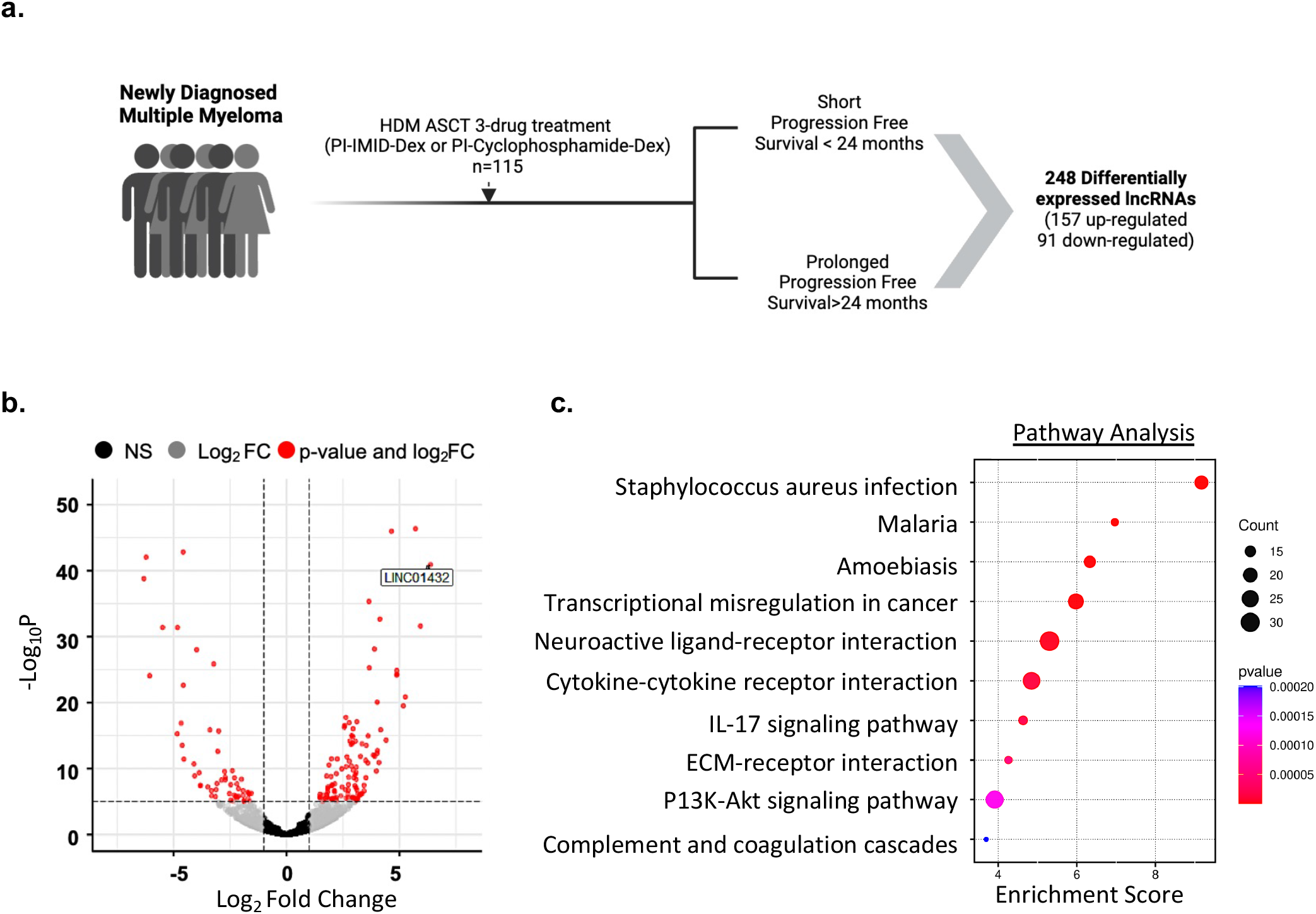
Identification of dysregulated lncRNAs associated with short progression-free survival to MM therapy. a. Schematic of the pipeline used to identify lncRNAs associated with short progression-free survival (PFS) to standard MM therapy. **b.** Identification of significantly differentially expressed lncRNAs in short PFS, as compared to prolonged PFS. **c.** Pathway analysis of differentially expressed genes associated with short PFS.

**Table 1:** Demographic details of NDMM patients separated into short and prolonged progression-free survival to standard multiple myeloma therapy.

| Characteristics | No. of Patients (%) |  | Total |
| --- | --- | --- | --- |
|  | Short PFS<br>(n=38) | Prolonged PFS<br>(n=77) | 115 |
| Age |  |  |  |
| ≤60 | 23 (33.3) | 46 (66.7) | 69 |
| ≥60 | 15 (32.6) | 31 (67.4) | 46 |
| Gender |  |  |  |
| Male | 22 (37.9) | 36 (62.1) | 58 |
| Female | 16 (28.1) | 41 (71.9) | 57 |
| Race |  |  |  |
| White | 31 (34.8) | 58 (65.2) | 89 |
| AA black | 4 (22.2) | 14 (77.8) | 18 |
| Other | 3 (37.5) | 5 (62.5) | 8 |
| Feature |  |  |  |
| Del17p | 1 (100) | 0 (0) | 1 |
| Amp1q | 17 (47.2) | 19 (52.8) | 36 |
| t4:14 | 5 (45.5) | 6 (54.5) | 11 |
| t8:14 | 10 (50.0) | 10 (50.0) | 20 |
| t11:14 | 4 (23.5) | 13 (76.5) | 17 |
| t14:16 | 1 (33.3) | 2 (66.7) | 3 |
| t14:20 | 0 (0) | 0 (0) | 0 |
| R-ISS |  |  |  |
| Stage I | 10 (24.3) | 31 (75.6) | 41 |
| Stage II | 23 (35.3) | 42 (64.6) | 65 |
| Stage III | 5 (55.5) | 4 (44.4) | 9 |

### LINC001432 is the top most significantly upregulated lncRNA in patients with short PFS

We focused our subsequent experimental analyses on characterizing the most significantly upregulated lncRNA in short PFS, as compared to prolonged PFS, termed *LINC01432 (*Fold change = 6.42, *p* = 6.11e^-43^), **Figure 1b** and **Figure 2a***. LINC01432* is a long intergenic non-protein coding RNA located on chromosome 20, has four exons, and is 693 nucleotides long. There is little-to-no current knowledge about *LINC01432*; it has only been reported to contain a SNP associated with male baldness in a single-trait genome-wide association study.^55^ Due to the heterogeneity and hyperploidy in several chromosomes observed in MM patients, we began by characterizing *LINC01432* by assessing different genetic subtypes of MM. We found that high expression of *LINC01432* was correlated with t(14;16) and Amp (1q) translocations (t[14;16] positive correlation = 0.57; Amp [1q] positive correlation = 0.14), **Supplementary** Figure 3a.

**Figure 2.**
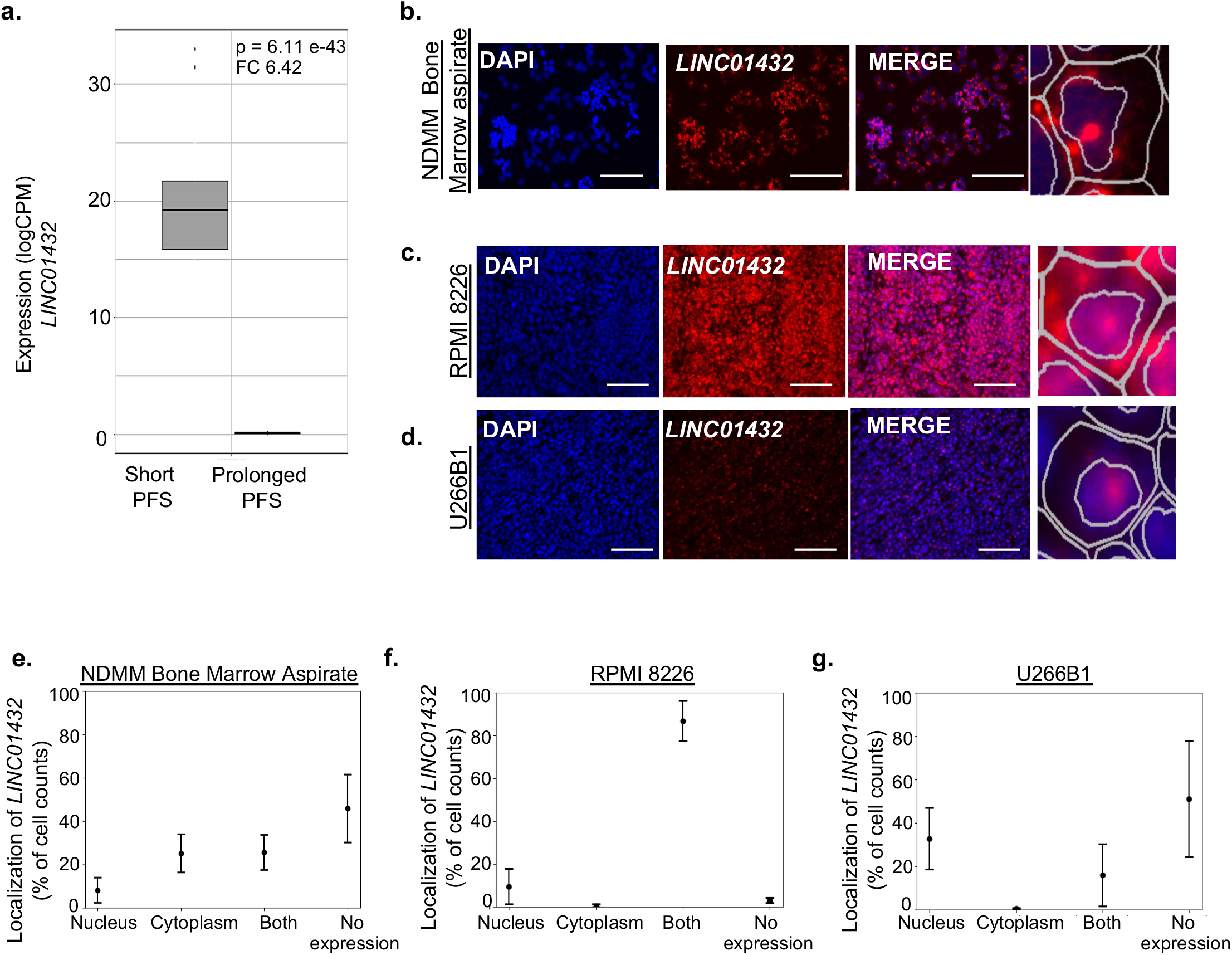
*LINC01432* is the most upregulated lncRNA in patients with short progression-free survival to MM therapy. a. Expression of *LINC01432* showing higher expression in short PFS compared to prolonged PFS. **b.** mFISH showing localized expression of *LINC01432* in newly diagnosed multiple myeloma patient bone marrow aspirates. **c**. mFISH showing localized expression of *LINC01432* in RPMI 8226 and **d.** U266B1 cell lines via subcutaneous injection in mouse. **e-g.** Quantification of nuclear and cytoplasmic co-localization (mFISH) of *LINC01432* in MM cell lines. A zoomed view of the cell is shown on each side of the panel. Scale bar = 20uM and 40 uM

To further characterize *LINC01432*, we analyzed its expression in a panel of MM cell lines and found that *LINC01432* is highly expressed in RPMI 8226 and OPM-2 cells, with low level expression detected in MM.1S, MM.1R, and U266B1 cells, **Supplementary** Figure 3b. Next, we confirmed the expression of *LINC01432* in NDMM bone marrow aspirates using mFISH, **Figure 2, b and e**. To assess the clinical significance of *LINC01432* in the context of MM, we subcutaneously injected mice with the MM cell lines RPMI 8226 and U266B1 to assess *in vivo* tumor growth and *LINC01432* expression. We found high expression of *LINC01432* in RPMI 8226 tumors and low expression in U266B1 tumors using mFISH, **Figure 2, c and d**. We determined that *LINC01432* is localized in both the cytoplasm and the nuclear compartments of RPMI 8226 cell line tumors, with 9.50% of cells exhibiting expression in the nucleus, 0.59% exhibiting expression in the cytoplasm, 86.91% exhibiting expression in both compartments, and 2.99% with no apparent *LINC01432* expression, **Figure 2f**. In the U266B1 cell line, which has low endogenous *LINC01432* expression levels, *LINC01432* expression was located in the nucleus in 32.79% of cells, in the cytoplasm of 0.44% of cells, in both compartments of 15.83% of cells, and expression was not detected in 50.94% of cells, **Figure 2g**. These data indicate that *LINC01432* is highly expressed in NDMM patient samples and in MM cell lines and is a novel lncRNA expressed in patients with short PFS to standard treatment.

### LINC01432 inhibits apoptosis and increases tumor growth

To investigate the molecular mechanisms through which *LINC01432* may induce a short PFS to standard MM therapy and to test its potential as a therapeutic target, we used CRISPR/Cas9 (CRISPR) to knockdown *LINC01432* expression in RPMI 8226 cells, **Figure 3a**, that have high endogenous expression levels, **Supplementary** Figure 3b. The development of a knockout cell line was unsuccessful due to a result of high cell death. Using the ApoTox-Glo Triplex Assay, we found that *LINC01432* knockdown significantly decreased viability (*p* = 0.03) and significantly increased apoptosis (*p* = 2.69e^-05^) in these cells, as compared to control CRISPR cells, **Figure 3b**. Increased apoptosis in *LINC01432* knockdown cells was further validated via Annexin V flow cytometry (*p* = 1.25e^-06^), **Figure 3c**. We then generated *LINC01432* overexpression from the U266B1 cell line, which have low endogenous *LINC01432* expression levels (**Figure 3d** and **Supplementary** Figure 3b) and found that these cells have significantly increased viability (*p* = 0.001) and significantly decreased apoptosis (ApoTox-Glo *p* = 0.04, AnnexinV *p* = 0.04), as compared to empty vector control cells, **Figure 3, e and f**. Next, we subcutaneously injected mice with *LINC01432* knockdown and control CRISPR cell lines and compared *in vivo* tumor growth **Supplementary** Figure 3c. This revealed significantly lower tumor volume in mice injected with *LINC01432* knockdown cells, as compared to control cells (Day 28 *p* = 0.04, Day 42 *p* = 0.02), **Figure 3g** **and Supplementary** Figure 3d. We similarly injected mice with *LINC01432* overexpression cells and empty vector controls and found that overexpression resulted in significantly higher tumor volume, as compared to controls (Day 14 *p* = 0.009, Day 21 *p* = 0.04, Day 28 *p* = 0.003, Day 35 *p* = 0.000), **Figure 3h** **and Supplementary** Figure 3e.

**Figure 3.**
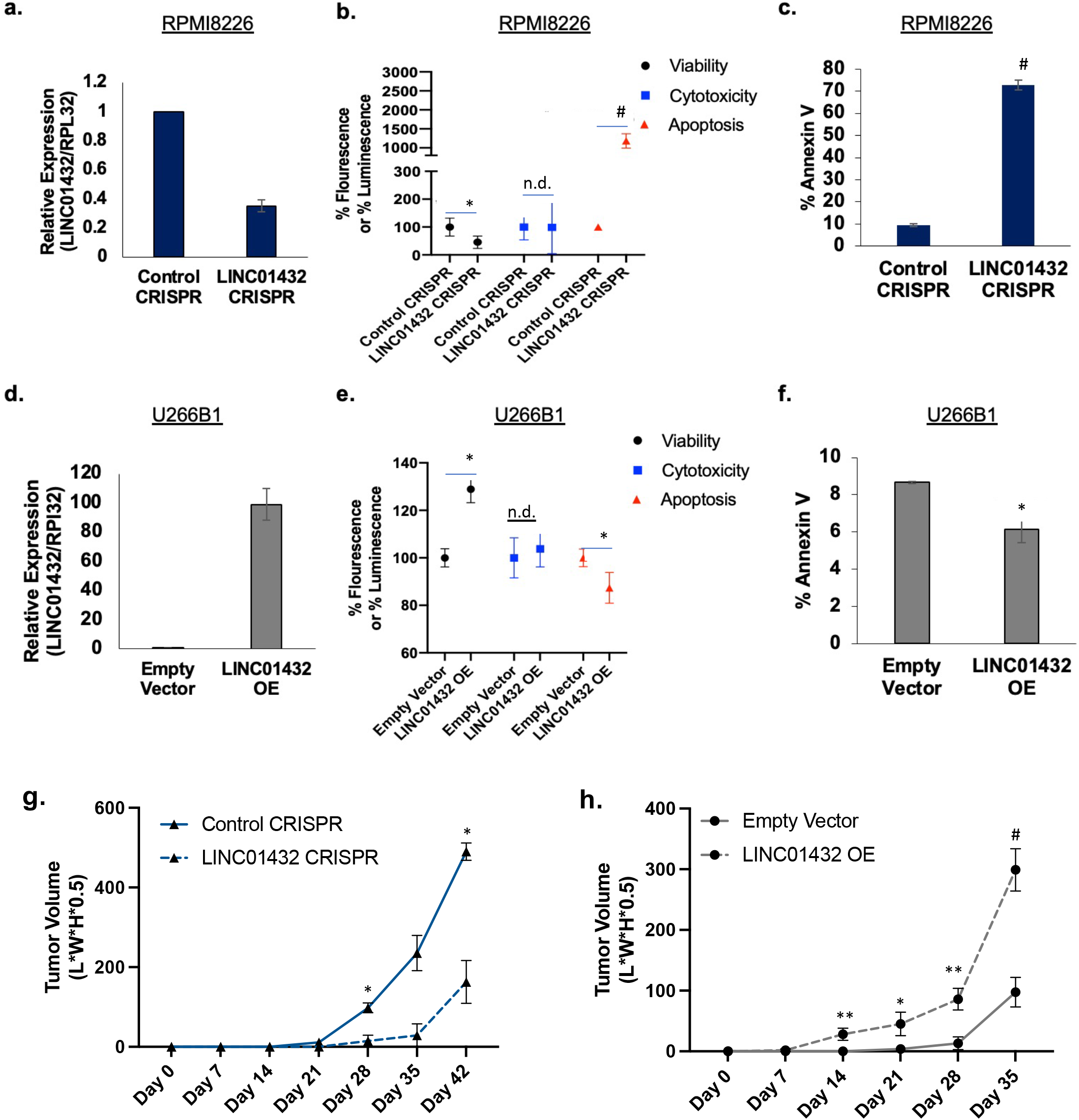
*LINC01432* promotes an aggressive phenotype. a. Expression of *LINC01432* in RPMI 8226 cells with CRISPR/Cas9-mediated *LINC01432* knockdown. **b.** *LINC01432* knockdown cells have decreased viability and increased apoptosis as measured via ApoTox-Glo assay, as compared to controls. **c.** Confirmation of increased apoptosis in *LINC01432* knockdown cells via Annexin V staining flow cytometry. **d.** Quantification of *LINC01432* overexpression U266B1 cells, as compared to empty vector controls. **e.** *LINC01432* overexpression cells have increased viability and decreased apoptosis as measured via ApoTox-Glo assay, as compared to controls. **f.** Confirmation of decreased apoptosis in *LINC01432* overexpression cells via Annexin V staining flow cytometry. **g.** Quantification of tumor growth following subcutaneous injection of *LINC01432* CRISPR/Cas9- mediated cell lines into NGS mice. **h.** Quantification of tumor growth following subcutaneous injection of *LINC01432* overexpression cell lines into NGS mice. *p < 0.05, **p < 0.005, #p < 0.0005

Next, we assessed the effects of *LINC01432* knockdown and overexpression on apoptotic markers, including *TP53* pathway genes, via RT-qPCR. We found that expression of apoptotic markers was significantly higher in RPMI 8226 *LINC01432* knockdown cells, as compared to controls (*TP53 p* = 0.004, *cMYC p* = 0.0002, *BAX p* = 5.96e^-13^). Similarly, we found that expression of apoptotic markers was significantly lower in tumors arising from *LINC01432* overexpression cells, as compared to controls (*TP53 p* = 2.04e^-09^, *cMYC p* = 1.57e^-10^, *BAX p* = 1.07e^-07^), **Supplementary** Figure 4**, a and b**. In addition, we detected a decrease in *γH2AX*, a marker of DNA double-stranded breaks, in tumors arising from *LINC01432* overexpression cells, **Supplementary** Figure 4c. These data provide evidence that *LINC01432* is highly expressed in MM cell lines and its expression is associated with increased viability and decreased apoptosis.

### LINC01432 binds to CELF2 protein

Many functional studies of lncRNAs, including our group’s research, have found that the ability of lncRNAs to bind with proteins and regulate downstream genes is integral to their roles in cancer and therapeutic resistance.^30,31^ As limited functional data on *LINC01432* is available, we utilized POSTAR3^47^ as a first step to identifying proteins which potentially bind to *LINC01432*. POSTAR3 is a unique, comprehensive database of post-transcriptional regulation and RNA-binding proteins that incorporates publicly available large-scale datasets on CLIP-sequencing, Ribo-sequencing, RNA secondary structure, and miRNA-mediated degradation events. This analysis identified that *LINC0143*2 to bind to the CELF2 RNA-binding protein from a publicly available CLIP-sequencing dataset ^56^, **Supplementary** Figure 5, **Figure 4, a and b**. Although the determined binding score was low (0.019), CELF2 (CUGBP Elav-like family) proteins are RNA-binding proteins with pleiotropic capabilities in RNA processing that have been found to compete with non-coding RNAs, including lncRNAs.^57^ CELF2 has been shown to bind lncRNAs to regulate downstream mRNAs, thereby promoting proliferation, migration, and tumor growth of multiple cancers,^58–62^ however, this has not yet been studied in MM. Thus, we investigated whether *LINC01432* binds to CELF2.

**Figure 4.**
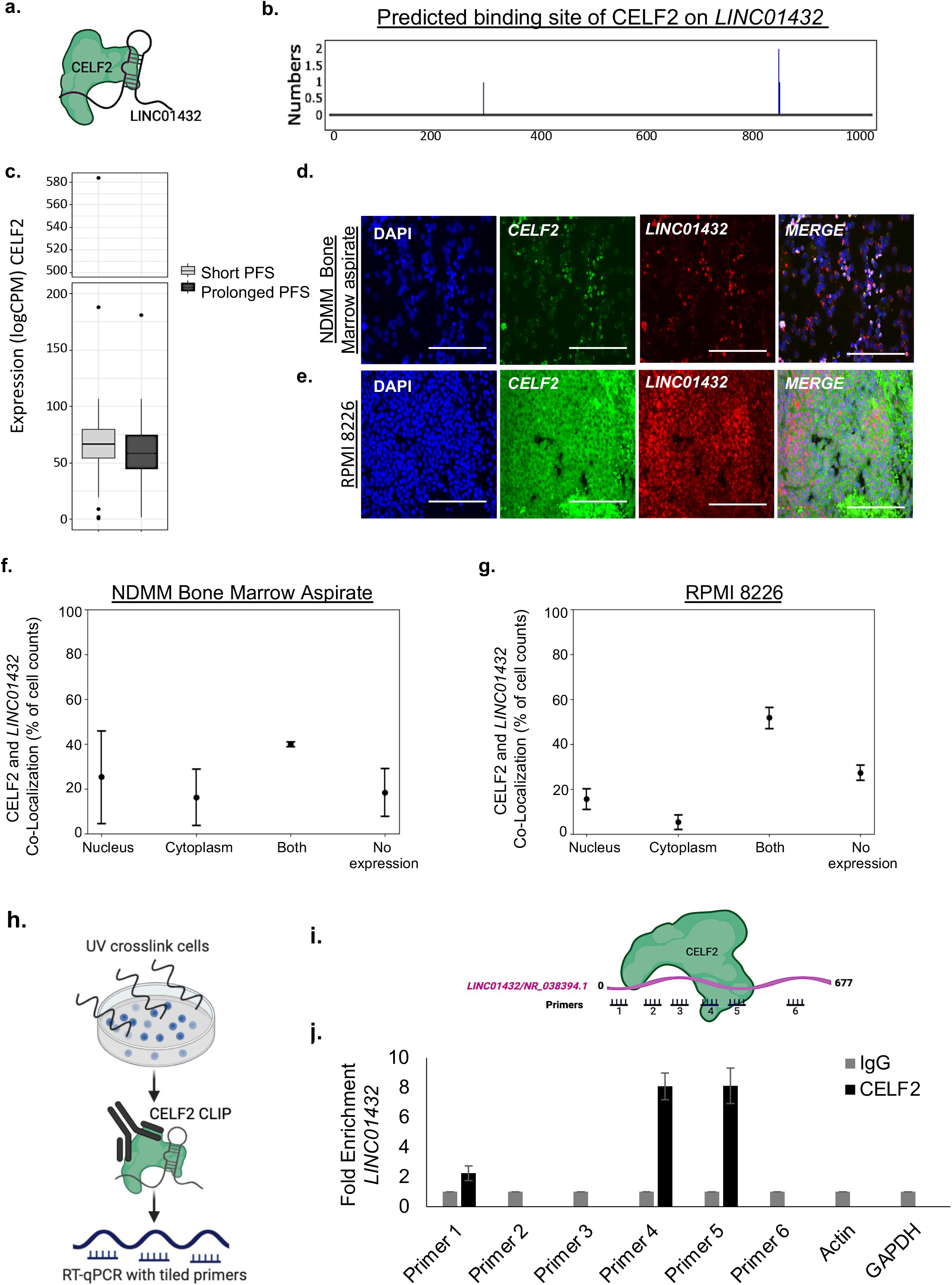
CELF2 binds to *LINC01432*. a. POSTAR3 prediction of CELF2 binding to *LINC01432* lncRNA. **b.** Predicted binding site of CELF2 on *LINC01432* by POSTAR3. **c**. Expression of CELF2 in short PFS compared to prolonged PFS, based on analysis of patient RNA sequencing data. **d. and e.** Localization of *LINC01432* lncRNA and CELF2 protein as determined by mFISH assay using *LINC01432* RNA probes combined with CELF2 antibodies using Immunohistochemistry in NDMM bone marrow aspirates and RPMI 8226 cells. **f. and g.** Quantification of nuclear and cytoplasmic localization of CELF2 and *LINC01432* in MM cells using QuPath**. h.** Schematic of *in vitro* iCLIP protocol**. i.** Schematic of tiled primers used to identify prospective CELF2 binding site on *LINC01432***. j.** CELF2 iCLIP RT-qPCR data, showing CELF2 binding to *LINC01432* compared to IgG negative control. Actin and GAPDH serve as negative gene controls. Scale bar = 40uM

Analysis of our NDMM patient RNA sequencing dataset revealed high level expression of CELF2 (logCPM > 50), but no significant differences in expression levels were identified between short PFS and prolonged PFS, **Figure 4c**. The Human Protein Atlas (proteinatlas.org) indicates that CELF2 expression is enriched in bone marrow (Tau score = 0.40) and localized to the nucleoplasm, vesicle, and midbody ring. Assessment of CELF2 expression in multiple blood cancer types indicated that CELF2 is highly expressed in leukemia, lymphoma, and MM (logCPM >10), **Supplementary** Figure 6a. Western blot analysis of CELF2 protein expression in whole cell lysates showed slight increased expression of CELF2 in both *LINC01432* RPMI 8226 knockdown (Fold Change = 1.41) and U266B1 overexpression cells (Fold Change = 1.58), as compared to controls, **Supplementary** Figure 6**, b and c.** mFISH analysis using *LINC01432* probes in combination with CELF2 protein immunohistochemistry in NDMM bone marrow aspirates indicated that CELF2 is expressed in both the nucleus and cytoplasm, **Figure 4, d and f**. Further, we found evidence of CELF2 and *LINC01432* co-localization in both cellular compartments in RPMI 8226 wild-type **Figure 4, e and g**, and U266B1 *LINC01432* overexpression cells, **Supplementary** Figure 7.

Next, we conducted iCLIP analysis to identify the regions of *LINC01432* that may be directly bound by CELF2, which combines UV cross-linking with immunoprecipitation and RT-qPCR to precisely map the binding sites of RNA-binding proteins, **Figure 4h**. RT-qPCR tiling primers spanning *LINC01432* showed direct binding of CELF2 to Tiling Primer 1 (Fold Change > 2), Tiling Primer 4 (Fold Change > 8), and Tiling Primer 5 (Fold Change > 8), as compared to IgG negative control, in RPMI 8226 cells with high level endogenous expression of *LINC01432*, **Figure 4, i and j**. These data indicate that *LINC01432* is bound by CELF2 protein in MM cell lines.

### Treating cells with LINC01432 locked nucleic acid antisense oligos (LNA ASOs) increases apoptosis

To assess the potential of *LINC01432* as a therapeutic target, we developed *LINC01432*-targeted LNA ASOs, which are increasingly being evaluated in clinical trials, along with control LNA ASOs.^63,64^ We treated RPMI 8226 cells and MM1.R cells with these LNA ASOs, **Figure 5a**, and confirmed knockdown of *LINC01432* expression, **Figure 5, b and d**. We next showed that LNA ASO-mediated *LINC01432* knockdown increased the proportion of cells in early apoptosis (MM1.R *p* = 0.003), late apoptosis (RPMI 8226 *p* = 0.005, MM1.R *p* = 0.005), and necrosis apoptosis (RPMI 8226 *p* = 0.013, MM1.R *p* = 0.02), as measured by flow cytometry, **Figure 5, c and d**. Further, we showed that LNA ASO-mediated CELF2 knockdown increased apoptosis in MM cell lines, **Supplementary** Figure 8**, a and b**. These data provide evidence that *LINC01432* inhibits apoptosis.

**Figure 5.**
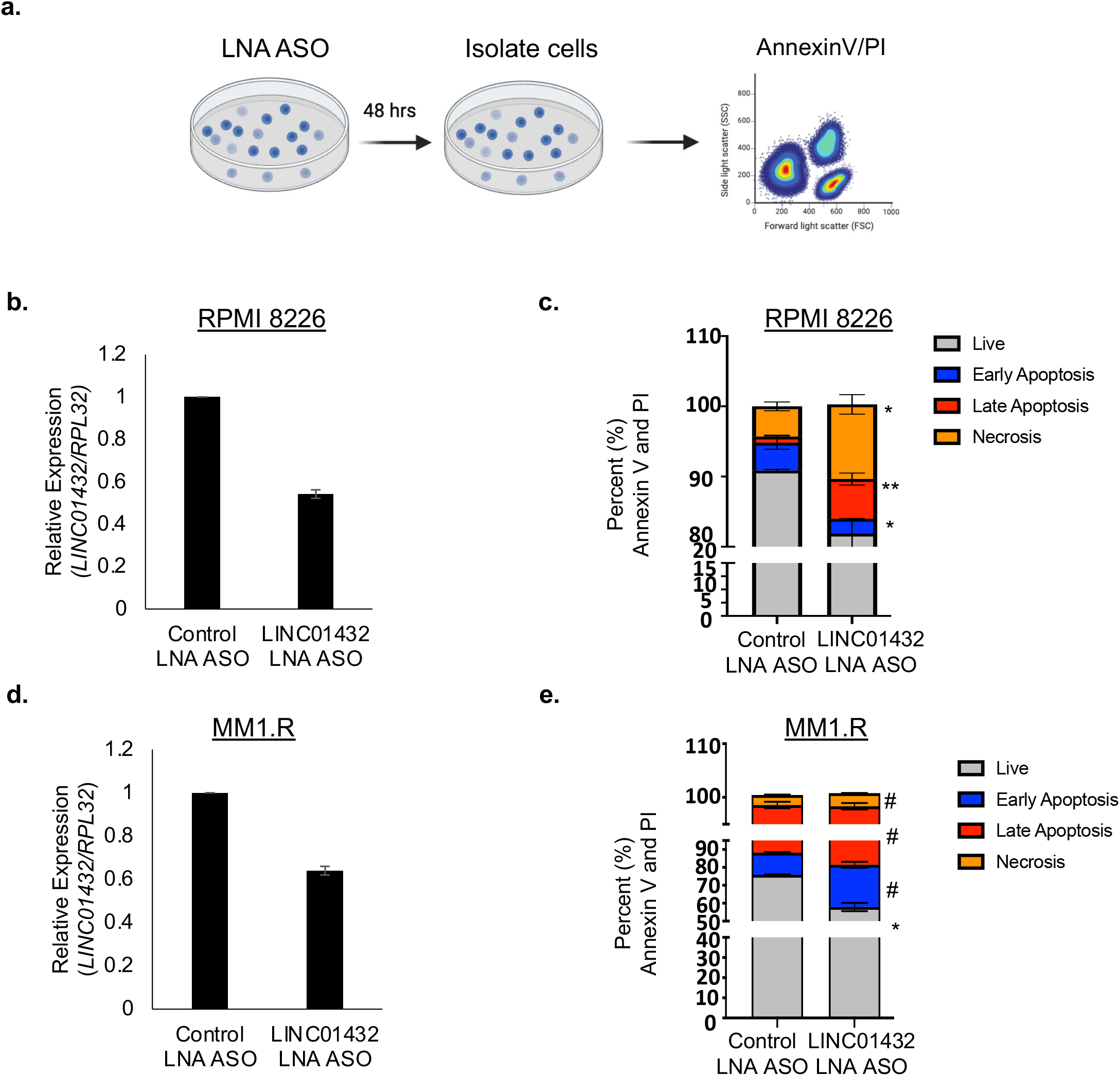
LNA ASO-mediated *LINC001432* knockdown induces apoptosis. a. Schematic of LNA ASO treatment for MM cells followed by flow cytometry. **b.** Expression of *LINC01432* in RPMI 8226 cells treated with Control LNA ASO or *LINC01432*-targeted LNA ASO, quantified by RT-qPCR. **c.** RPMI 8226 cells with LNA ASO- mediated LINC001432 knockdown on apoptosis, as measured via Annexin V and PI flow cytometry. **b.** Expression of *LINC01432* in MM1.R cells treated with Control LNA ASO or *LINC01432*-targeted LNA ASO, quantified by RT-qPCR. **c.** MM1.R cells with LNA ASO-mediated LINC001432 knockdown on apoptosis, as measured via Annexin V and PI flow cytometry *p < 0.05, **p < 0.005, #p < 0.0005

In summary, our study identified differentially expressed lncRNAs associated with patients who had short PFS to standard MM therapy and determined that *LINC01432* bound to CELF2 and that LNA knockdown in MM cells promotes apoptosis and decreases viability suggesting the role of *LINC01432* and the *LINC01432*-CELF2 complex in promoting PFS to the standard treatment, **Figure 6**.

**Figure 6.**
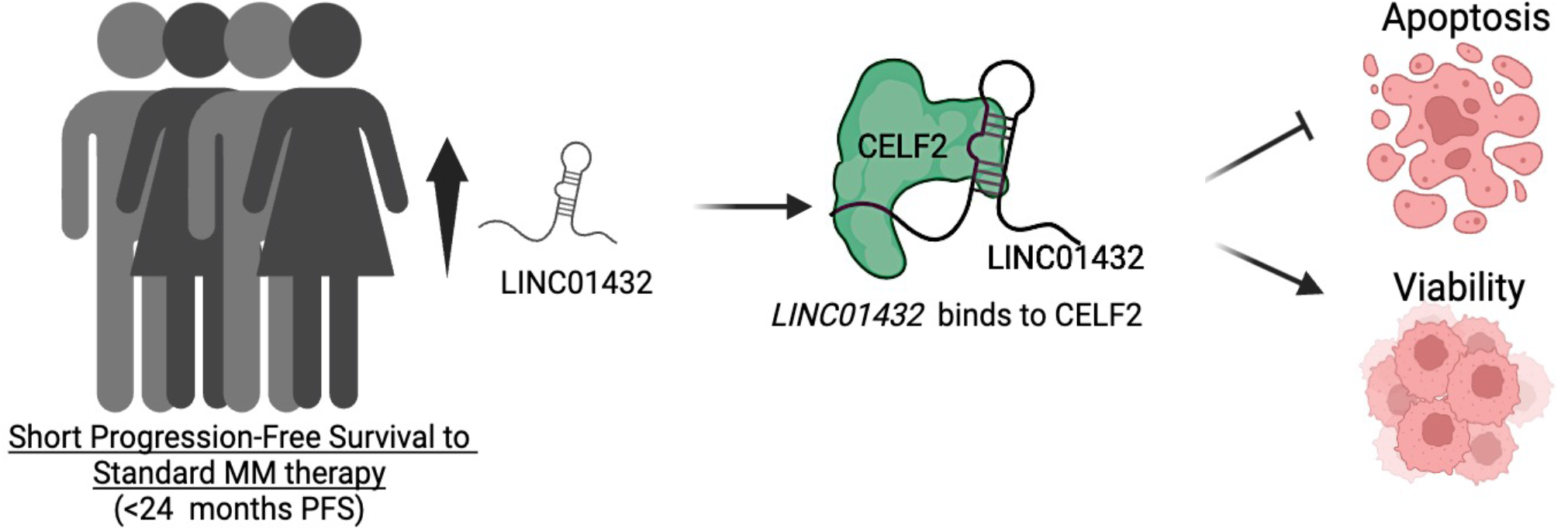
Schematic of overall outcomes. *LINC01432* is highly expressed in newly diagnosed multiple myeloma patients, is bound by CELF2 protein, and together they inhibit apoptosis and promote cell viability. Created with Biorender.com

## Discussion

The development of drug resistance results in most patients exhausting all available treatment options and relapsing.^6,65^ While some advances in MM treatment are emerging through clinical trials of novel cellular immunotherapies targeting immune cells, including chimeric antigen receptor T cell therapies,^66–69^unfortunately, interpatient heterogeneity has hindered the elucidation of the molecular mechanisms that control the progression of plasma cells in patients with MM.^27,70–72^ Thus, knowledge surrounding mechanisms and biomarkers related to treatment resistance in MM patients are critical for novel therapy development.

In this study, we identified differentially expressed lncRNAs in NDMM patients who exhibited a short PFS to the standard MM therapy. The most upregulated annotated lncRNA, *LINC01432*, was found to bind to the CELF2 protein, leading to inhibition of apoptosis and promotion of cell viability. CELF2 has been previously reported to bind to lncRNAs and regulate downstream mRNAs, thereby promoting proliferation, migration, and tumor growth in multiple forms of cancer,^60–62,73–80^ however, this has not yet been studied in the context of NDMM. Interestingly, CELF2 expression patterns vary in different developmental and differentiation stages.^57^ In cancer, CELF2 is found to be localized to the nucleus, where it is associated with alternative splicing and transcript editing, in RNA granules, where it regulates mRNA stability, and in the cytoplasm, where it regulates pre-miRNA maturation, translation, and alternative polyadenylation.^56,81–84^ Here, we determined that CELF2 shows different patterns of expression in MM cell lines with differential levels of endogenous *LINC01432* expression. In the presence of *LINC01432,* CELF2 was localized to the cytoplasm. We observed that co-localization allowed the binding of *LINC01432* to CELF2 to inhibit apoptosis, although more studies need to be conducted to determine if this is a result of the specific interaction or is solely dependent on the increased expression of *LINC01432*. In addition, POSTAR3 also predicted *LINC01432* binding to the protein AGO2, thereby we believe that there may be additional proteins that may bind to *LINC01432,* as lncRNAs are known to interact with multiple RNA binding proteins. ^85,86^ As we are in the earliest stages of understanding *LINC01432* tumor biology, this study allows us to predict that *LINC01432-*CELF2 interaction may play a larger role in the pathogenesis of MM. Future studies are needed to better understand this interaction and other potential protein interactions in MM and to fully characterize its role in the development of resistance to chemotherapy.

One promising aspect of lncRNAs that makes them ideal novel targets for the development of RNA therapeutics is their tissue and cell specific expression patterns ^87,88^. ASOs are an emerging class of RNA-based therapeutic drugs that can be easily modified and optimized for clinical development.^89–92^ To date, there are 128 registered clinical trials of ASOs for the treatment of several diseases, including cancer.^93^ ASOs have been shown to be a powerful tool for therapeutically targeting lncRNAs.^94–97^ We developed a *LINC01432*-targeted LNA ASO and demonstrated its potential use to treat *LINC01432*-mediated decreased apoptosis and increased viability in MM cell lines. Further, treating MM cell lines with *LINC01432*-targeted LNA ASOs showed an increased apoptosis *in vitro*. This study represents a preliminary investigation into the use of LNA ASO to downregulate *LINC01432* lncRNA; future studies are being conducted to provide evidence of its clinical significance. Many unanswered questions remain regarding the exact mechanism through which *LINC01432* regulates downstream pathways and mediates epigenetic regulation while bound to CELF2. In conclusion, our study provides preliminary insights into the role of lncRNA expression in NDMM patients who exhibit a short PFS to standard MM therapy and identifies a novel potential target for the development of future MM therapies.

## Acknowledgments

J.S.F. received funding from the Faculty Diversity Scholar Award from the Washington University Department of Medicine and the Longer Life Foundation. We would like to thank Dr. John DiPersio for cell lines, support, and guidance for this project. We thank the Alvin J. Siteman Cancer Center at Washington University School of Medicine and Barnes-Jewish Hospital in St. Louis, MO, for utilizing the Siteman Flow Cytometry core that provided flow cytometry service. We also thank the Genome Engineering and Stem Cell Center for help in developing CRISPR cell lines. We appreciate the help and expertise from Washington University in St. Louis Multiple Myeloma Tissue Banking Protocol for guidance on sample assessment for sequencing data and access to myeloma tissue samples. We thank the Alvin J. Siteman Cancer Center at Washington University School of Medicine and Barnes-Jewish Hospital in St. Louis, MO., for using the Siteman Cancer Center Tissue Procurement Core, which provides cell sorting service. The Siteman Cancer Center is partly supported by an NCI Cancer Center Support Grant #P30 CA091842.The development of this manuscript was supported by the Scientific Editing Service of the Institute of Clinical and Translational Sciences at Washington University in St. Louis, funded by grant UL1TR002345 from the National Center for Advancing Translational Sciences. The content is solely the responsibility of the authors and does not necessarily represent the official view of the NIH. This work is also supported by NCI R35 Outstanding Investigator Award; R35 CA210084 (JD) and the Riney Blood Cancer Research Fund.

## Authorship Contributions

R.M. wrote and edited the manuscript, conducted *in vitro* assays, mouse experiments, mFISH, and analyzed data. P.T. edited manuscript, developed cell lines and conducted in vitro assays. A.P. edited manuscript, conducted in vitro assays, and western blots. D.D. edited manuscript, analyzed and processed RNA sequencing data, and mFISH images, K.B. analyzed and processed RNA sequencing data, S.G. conducted and analyzed *in vitro* assays and mFISH, V.B. conducted and analyzed *in vitro* assays, S.D., M.A., C.B., Y.R., J.K. conducted *in vitro* assays, Y.J. and K.W. analyzed mFISH, R.J., M.F., J.F. provided guidance on analysis of RNA sequencing data cohorts, R.J. provided guidance and analysis of sequencing data, M.S., L.D., R.V., and J.D. provided project guidance. J.D. provided MM cell lines. R.V. provided samples. J.S-F. conducted *in vitro* assays, and mouse experiments, designed and directed experimental studies, and wrote a manuscript, which all authors reviewed.

### Disclosure of Conflicts of Interest Support

M.S. (Consulting for DSMB and endpoint adjudication work: Sorrento, Marker Therapeutics, GSK, Kura Oncology, and NovoNordisk, Research support: Incyte, Janssen, Takeda, Ichnos Biosciences, Fate Therapeutics, and Karyopharm), R.V. (Consulting: BMS, Sanofi, Janssen, Karyopharm, Pfizer, Regenron, Research: BMS, Sanofi, Takeda), and J.D (Consulting: Rivervest, Bluebird Bio, Vertex, HcBiosciences, SPARC), Equitiy-ownership WUGEN and Magenta, Research support: Macrogenics, Bioline, Incyte.

**Correspondence** and requests for materials should be addressed to Jessica M. Silva-Fisher.

---

**Supplementary Figure 1:**
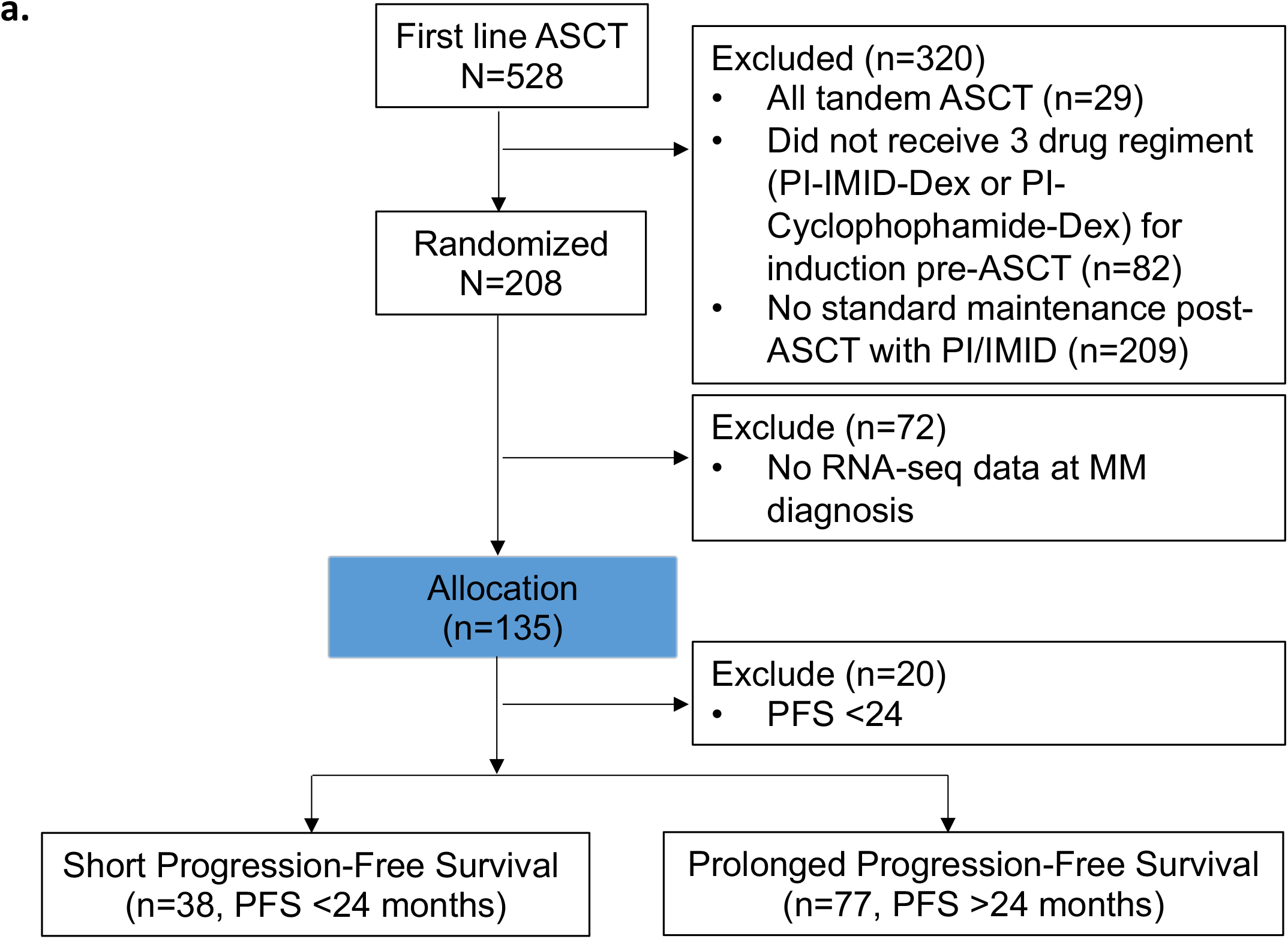
Identification of newly diagnosed multiple myeloma patients with short progression-free survival to standard therapy a. Pipeline to identify newly diagnosed multiple myeloma patients samples with short or prolonged progression-free survival to therapy.

**Supplementary Figure 2:**
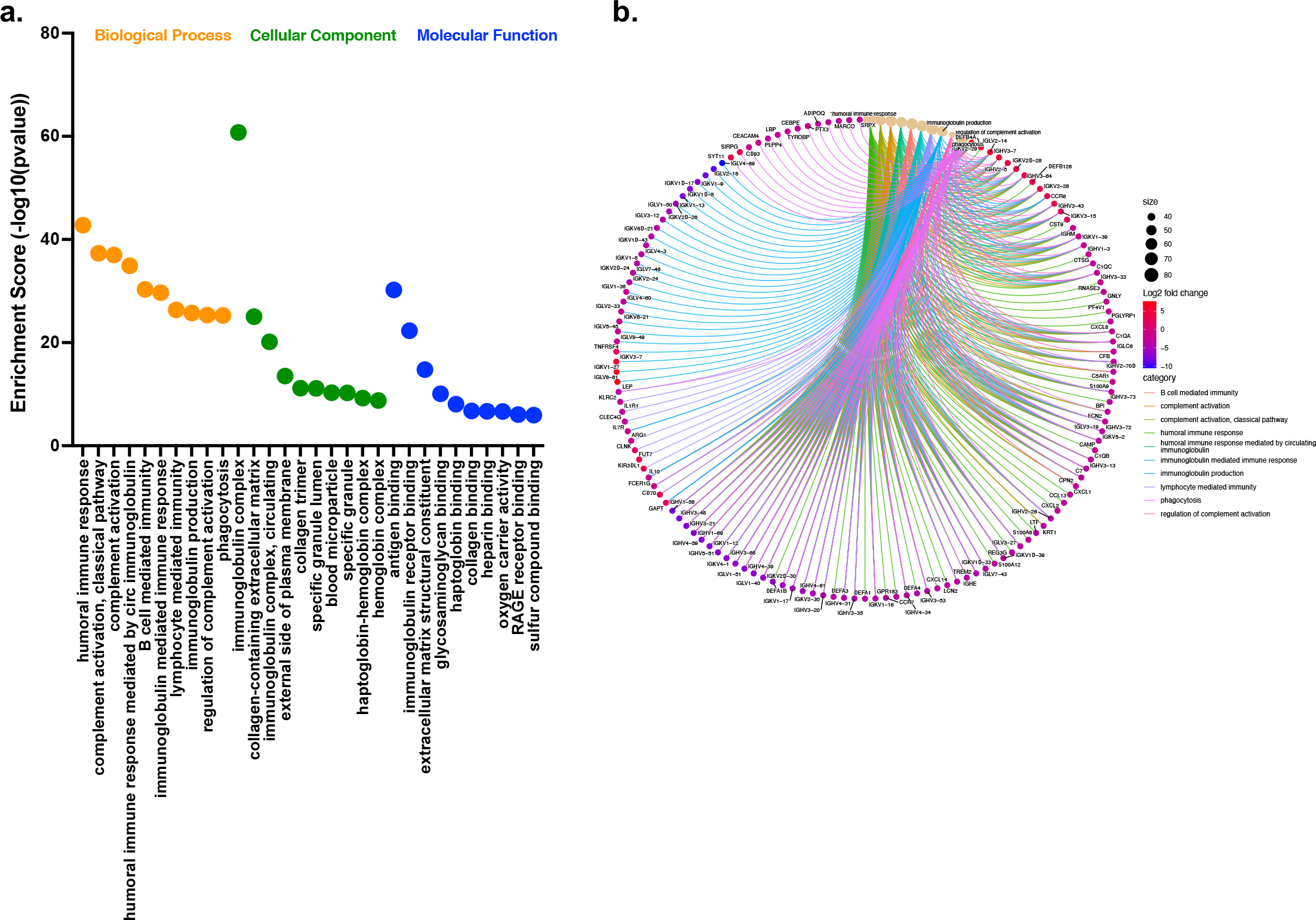
GO analysis comparing short to prolonged progression-free survival to MM therapy **a. Gene ontology and b. pathway analysis of genes associated with short PFS.**

**Supplementary Figure 3:**
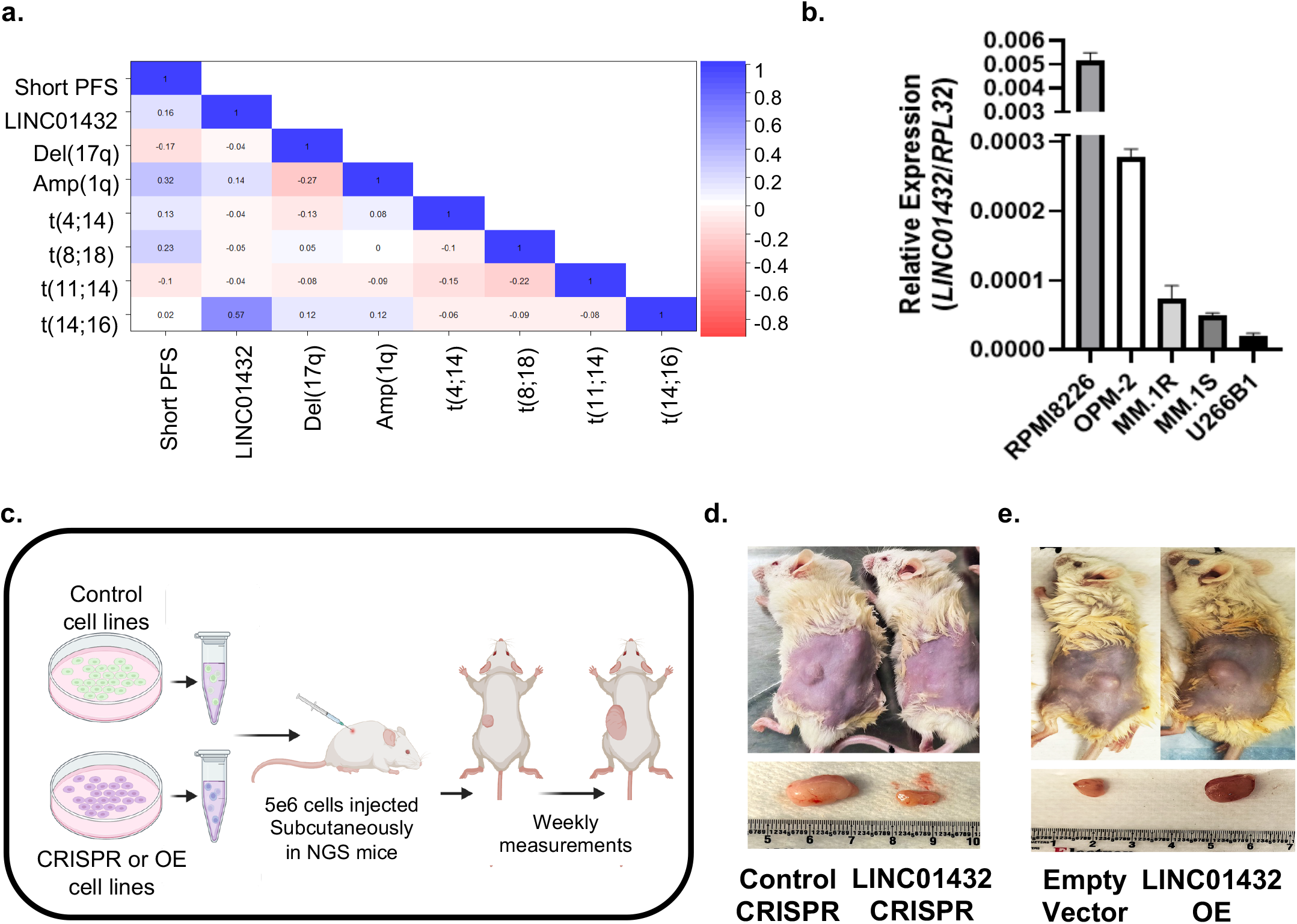
*LINC01432* expression and tumor growth **a.** Expression correlation of short progression-free survival (PFS) genes, *LINC01432*, and known multiple myeloma translocations **b.** Expression of *LINC01432* by RT-qPCR in myeloma cell line panel **c.** Schematic of subcutaneous injection of cell lines into NGS mice. **d.** Representative images of *LINC01432* knockdown cell-induced tumor growth sizes at Day 42 following subcutaneous injection into NGS mice, compared to control CRISPR cell-induced tumors. **e.** Representative images of *LINC01432* overexpression cell-induced tumor growth sizes at Day 35 following subcutaneous injection into NGS mice, compared to empty vector control cell-induced tumors.

**Supplementary Figure 4:**
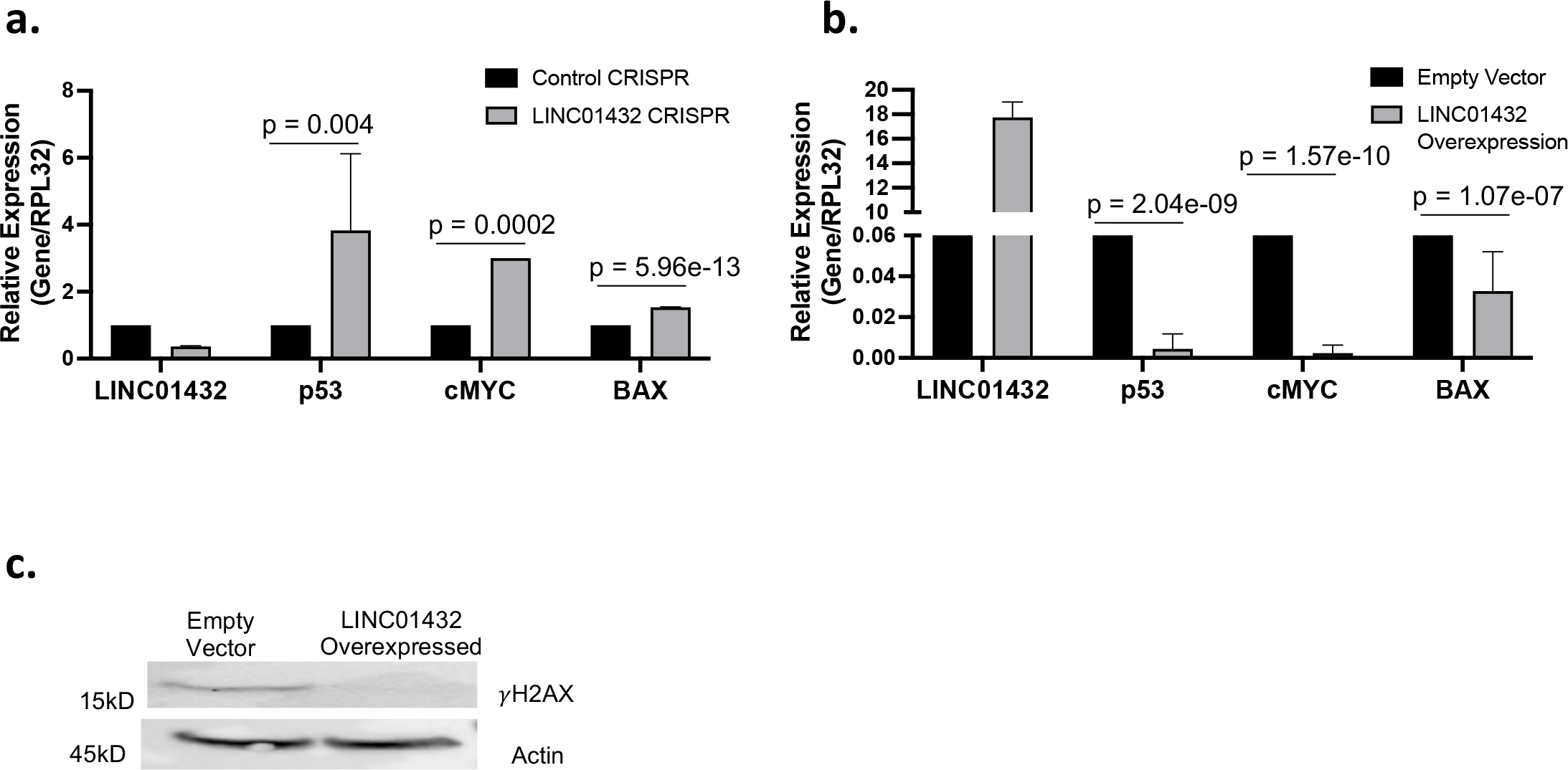
*LINC01432* inhibits apoptosis TP53 pathway **a.** Expression of TP53 apoptotic pathway genes in *LINC01432* knockdown cell lines or **b.** overexpression cell lines. **c.** Western blot showing a decrease in yH2AX in *LINC01432* overexpression cell lines.

**Supplementary Figure 5:**
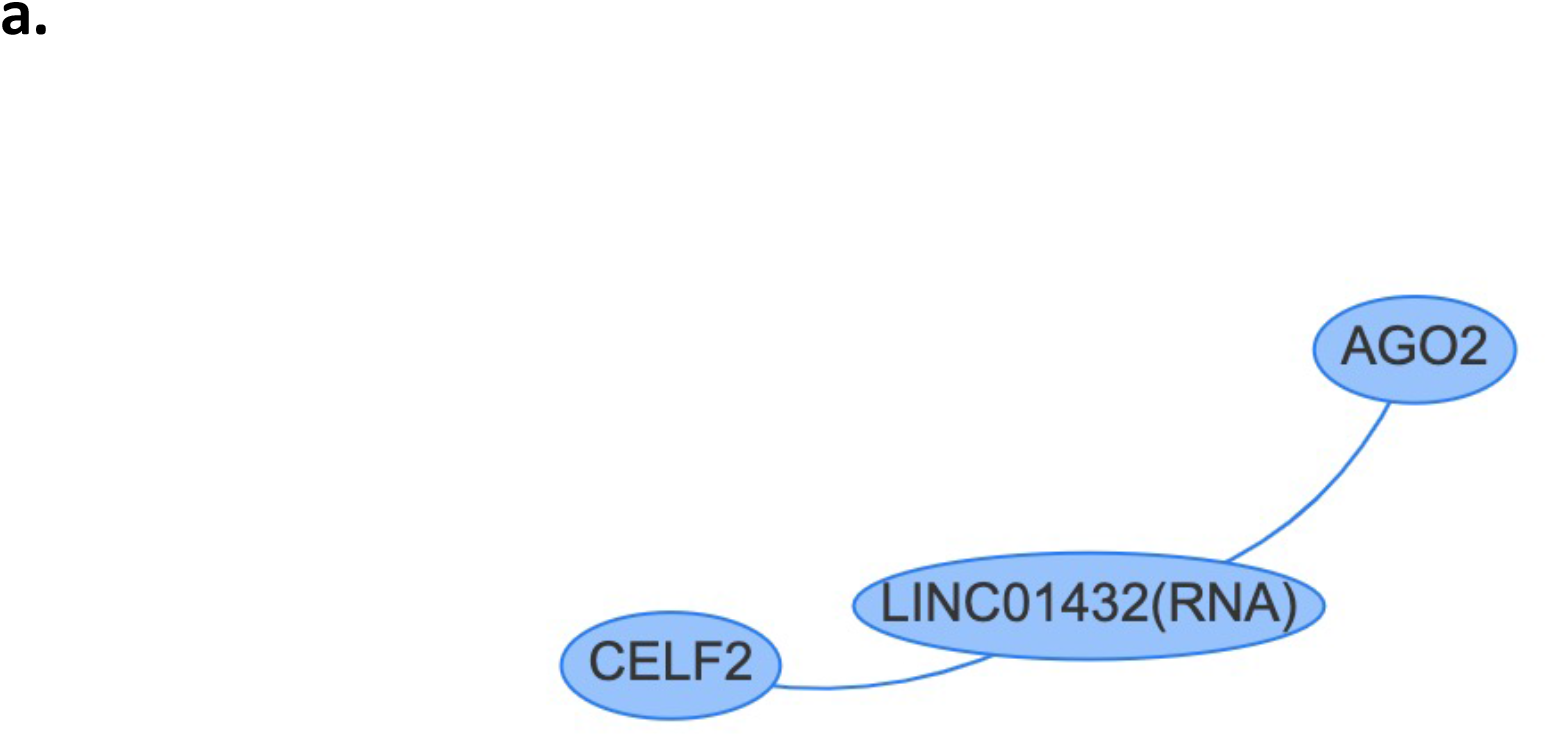
POSTAR3 RNA binding site prediction for ***LINC01432 a.*** *LINC01432* interaction network displayed using POSTAR3 program identified binding to CELF2 and AGO2 proteins.

**Supplementary Figure 6:**
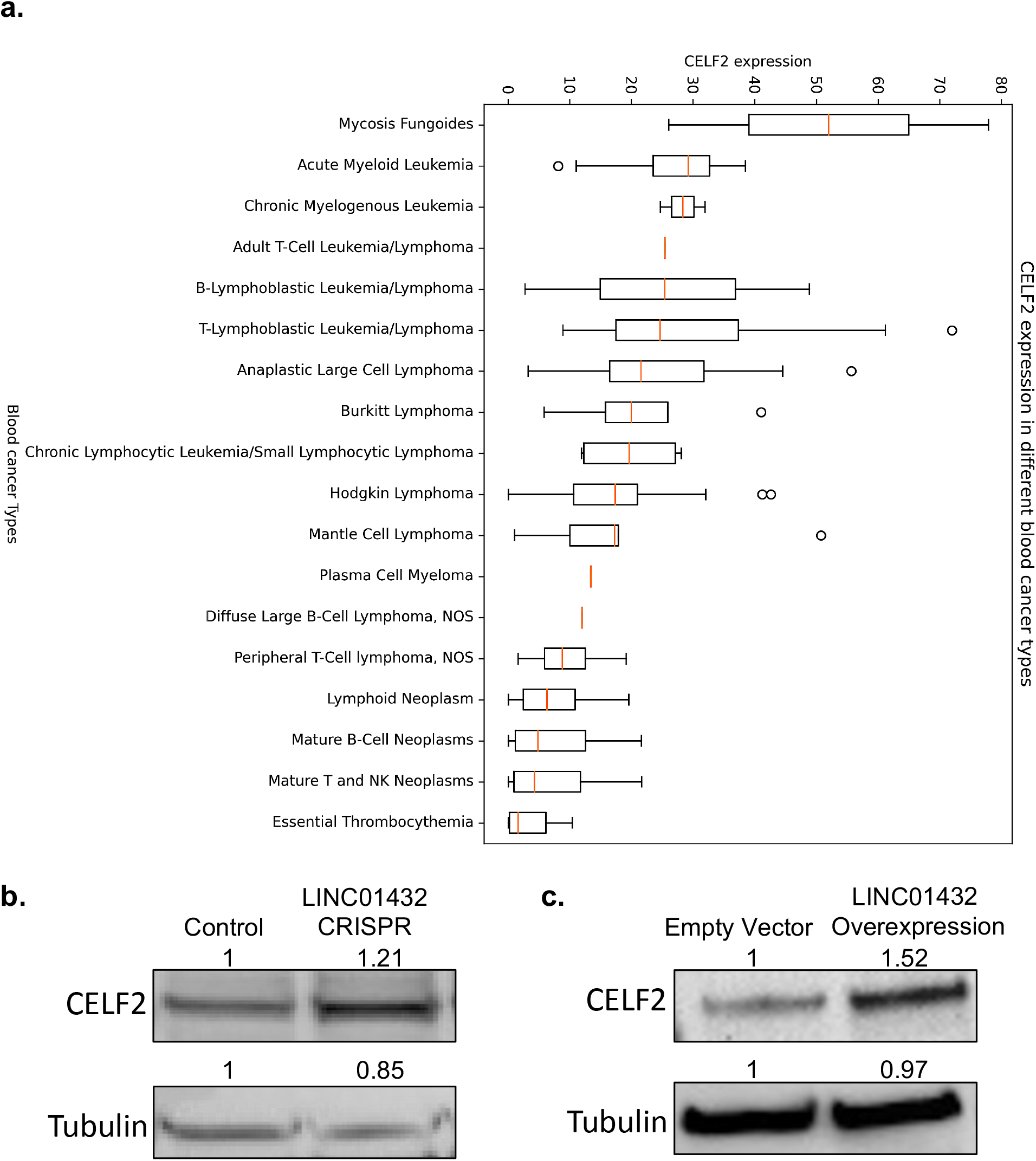
CELF2 RNA protein binding in multiple myeloma **a.** Expression of *CELF2* in blood cancers from Cancer Cell Line Encyclopedia **b.** CELF2 protein expression from *LINC01432* CRISPR cell lines or **c.** overexpression cell lines.

**Supplementary Figure 7:**
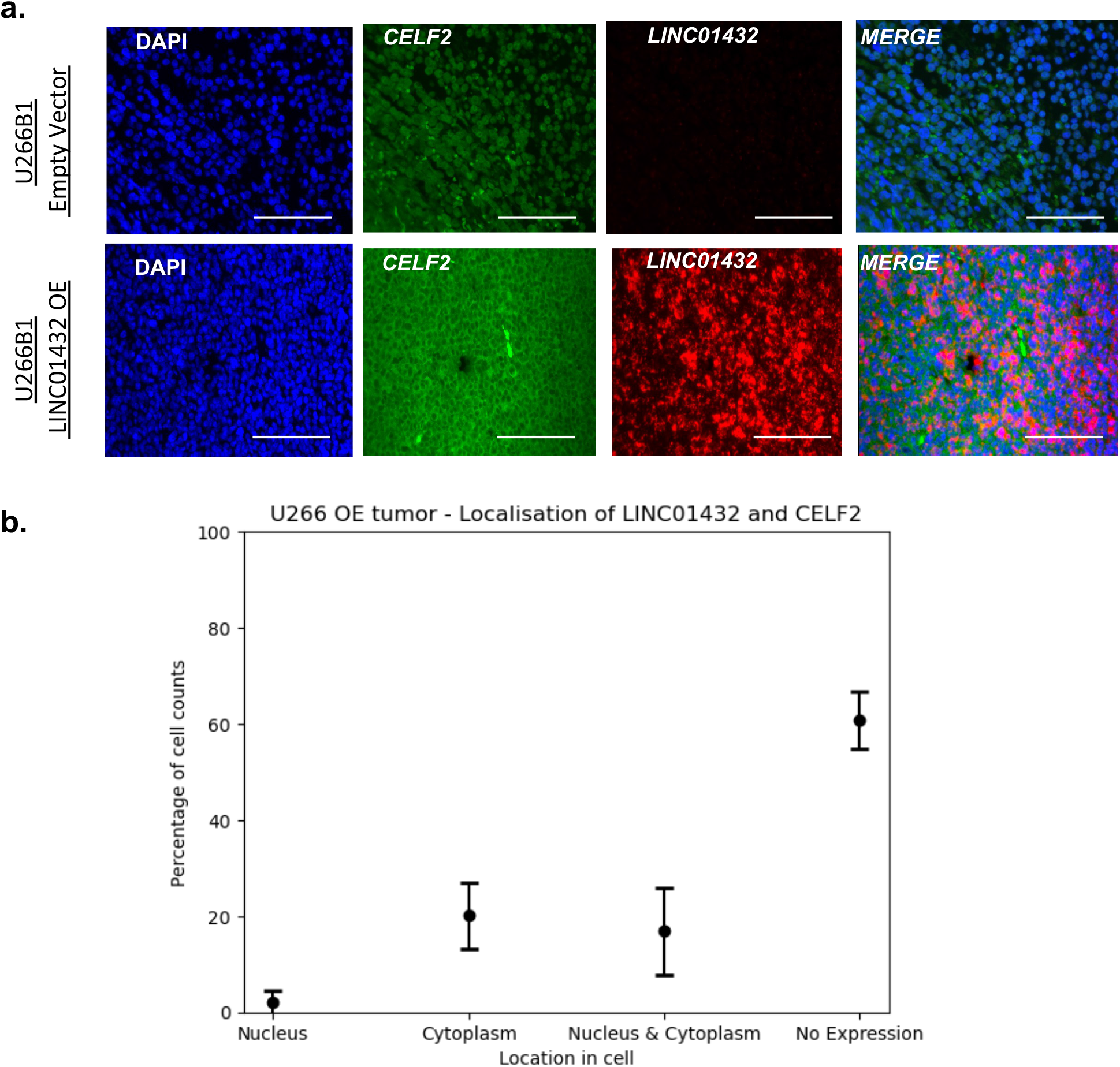
LINC01432 binds to CELF2 RNA protein binding in over expression cells **a.** Localization of *LINC01432* lncRNA and CELF2 protein as determined by mFISH assay using *LINC01432* RNA probes combined with CELF2 antibodies using Immunohistochemistry in U266B1 empty vector control cells, and *LINC01432* overexpression cells. **b.** Quantification of nuclear and cytoplasmic localization of CELF2 and *LINC01432* in U266B1 empty vector control cells, and *LINC01432* overexpression cells using QuPath.

**Supplementary Figure 8:**
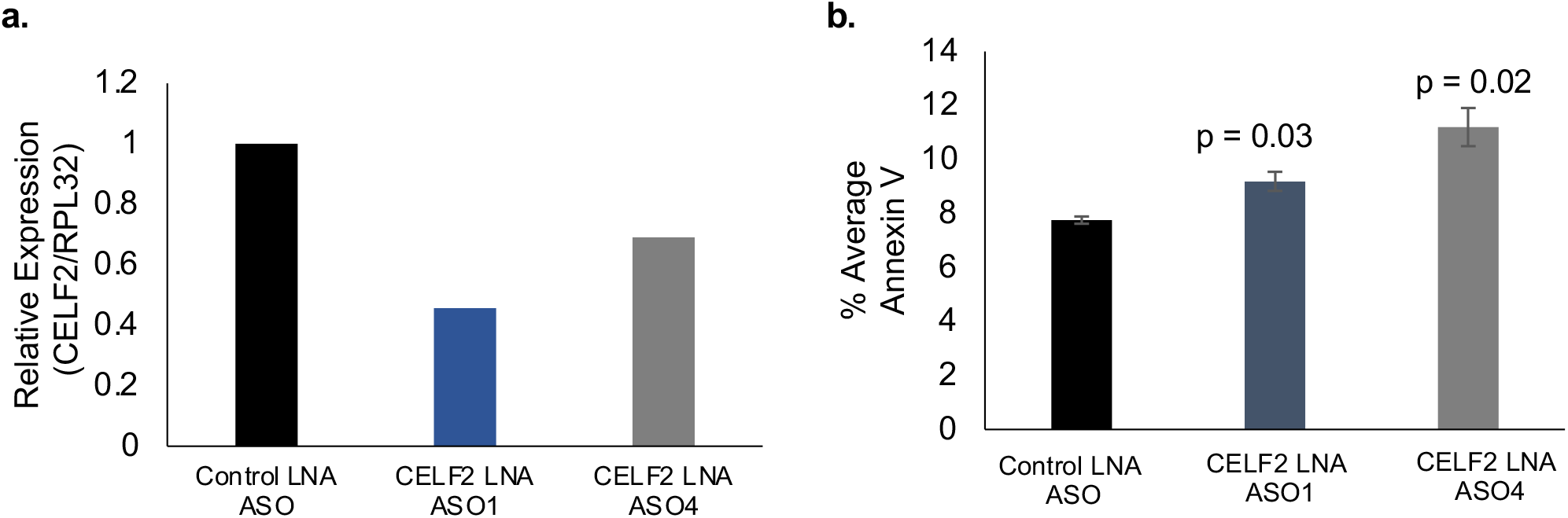
CELF2 induces apoptosis **a.** Expression of *CELF2* in U266B1 cells treated with control locked nucleic acid oligonucleotides (LNA ASO), or CELF2 LNA ASO1, or CELF2 LNA ASO4 **b.** CELF2 LNA ASO-treated cells induce apoptosis.

## References

1. Siegel, R.L., Giaquinto, A.N. & Jemal, A. Cancer statistics, 2024. CA: a cancer journal for clinicians 74, 12–49 (2024).

2. Samuels, B.L. & Bitran, J.D. High-dose intravenous melphalan: a review. Journal of clinical oncology : official journal of the American Society of Clinical Oncology 13, 1786–1799 (1995).

3. Falco, P., et al. Melphalan and its role in the management of patients with multiple myeloma. Expert Rev Anticancer Ther 7, 945–957 (2007).

4. Martino, M., et al. Addressing the questions of tomorrow: melphalan and new combinations as conditioning regimens before autologous hematopoietic progenitor cell transplantation in multiple myeloma. Expert Opin Investig Drugs 22, 619–634 (2013).

5. Minnie, S.A. & Hill, G.R. Immunotherapy of multiple myeloma. J Clin Invest 130, 1565–1575 (2020).

6. Gajek, A., et al. Chemical modification of melphalan as a key to improving treatment of haematological malignancies. Scientific reports 10, 4479 (2020).

7. Frede, J., et al. Dynamic transcriptional reprogramming leads to immunotherapeutic vulnerabilities in myeloma. Nature cell biology 23, 1199–1211 (2021).

8. Silva, J. & Smith, D. Long non-coding RNAs and Cancer (Caister Academic Press, La Jolla, California, 2012).

9. Cabanski, C.R., et al. Pan-cancer transcriptome analysis reveals long noncoding RNAs with conserved function. RNA Biol 12, 628–642 (2015).

10. White, N.M., et al. Transcriptome sequencing reveals altered long intergenic non-coding RNAs in lung cancer. Genome Biol 15, 429 (2014).

11. Qian, Y., Shi, L. & Luo, Z. Long Non-coding RNAs in Cancer: Implications for Diagnosis, Prognosis, and Therapy. Front Med (Lausanne) 7, 612393 (2020).

12. Saltarella, I., et al. The Landscape of lncRNAs in Multiple Myeloma: Implications in the "Hallmarks of Cancer", Clinical Perspectives and Therapeutic Opportunities. Cancers (Basel) 14(2022).

13. Coira, I.F., Rincon, R. & Cuendet, M. The Multiple Myeloma Landscape: Epigenetics and Non-Coding RNAs. Cancers (Basel) 14(2022).

14. Leng, S., Qu, H., Lv, X. & Liu, X. Role of ncRNA in multiple myeloma. Biomark Med 16, 1181–1191 (2022).

15. Yang, C., et al. Long non-coding RNAs in multiple myeloma (Review). Int J Oncol 62(2023).

16. Zeng, T., et al. Identification and validation of a cellular senescence-related lncRNA signature for prognostic prediction in patients with multiple myeloma. Cell Cycle 22, 1434–1449 (2023).

17. Garitano-Trojaola, A., Agirre, X., Prosper, F. & Fortes, P. Long non-coding RNAs in haematological malignancies. International journal of molecular sciences 14, 15386–15422 (2013).

18. Carrasco-Leon, A., Amundarain, A., Gomez-Echarte, N., Prosper, F. & Agirre, X. The Role of lncRNAs in the Pathobiology and Clinical Behavior of Multiple Myeloma. Cancers (Basel) 13(2021).

19. Shen, X., et al. Knockdown of long non-coding RNA PCAT-1 inhibits myeloma cell growth and drug resistance via p38 and JNK MAPK pathways. J Cancer 10, 6502–6510 (2019).

20. Wang, Y., et al. Long noncoding RNA H19 promotes vincristine resistance in multiple myeloma by targeting Akt. Cell Mol Biol (Noisy-le-grand) 66, 76–80 (2020).

21. Yang, L.H., et al. LncRNA ANRIL promotes multiple myeloma progression and bortezomib resistance by EZH2-mediated epigenetically silencing of PTEN. Neoplasma 68, 788–797 (2021).

22. Enukashvily, N.I., et al. Pericentromeric Non-Coding DNA Transcription Is Associated with Niche Impairment in Patients with Ineffective or Partially Effective Multiple Myeloma Treatment. International journal of molecular sciences 23(2022).

23. Zhou, F. & Guo, L. Lncrna ANGPTL1-3 and its target microRNA-30a exhibit potency as biomarkers for bortezomib response and prognosis in multiple myeloma patients. Hematology 27, 596–602 (2022).

24. Zang, X., et al. LncRNA MEG3 promotes the sensitivity of bortezomib by inhibiting autophagy in multiple myeloma. Leuk Res 123, 106967 (2022).

25. Ren, Y., et al. Expression of NEAT1 can be used as a predictor for Dex resistance in multiple myeloma patients. BMC cancer 23, 630 (2023).

26. Butova, R., Vychytilova-Faltejskova, P., Souckova, A., Sevcikova, S. & Hajek, R. Long Non-Coding RNAs in Multiple Myeloma. Non-coding RNA 5(2019).

27. Meng, H., Han, L., Hong, C., Ding, J. & Huang, Q. Aberrant lncRNA Expression in Multiple Myeloma. Oncol Res 26, 809-816 (2018).

28. Carrasco-Leon, A., et al. Characterization of complete lncRNAs transcriptome reveals the functional and clinical impact of lncRNAs in multiple myeloma. Leukemia 35, 1438–1450 (2021).

29. Wu, Z. & Lin, Y. Long noncoding RNA LINC00515 promotes cell proliferation and inhibits apoptosis by sponging miR-16 and activating PRMT5 expression in human glioma. OncoTargets and therapy 12, 2595–2604 (2019).

30. Silva-Fisher, J.M., et al. Long non-coding RNA RAMS11 promotes metastatic colorectal cancer progression. Nature communications 11, 2156 (2020).

31. Eteleeb, A.M., et al. LINC00355 regulates p27(KIP) expression by binding to MENIN to induce proliferation in late-stage relapse breast cancer. NPJ Breast Cancer 8, 49 (2022).

32. Li, S., et al. exoRBase: a database of circRNA, lncRNA and mRNA in human blood exosomes. Nucleic Acids Res 46, D106–D112 (2018).

33. Wu, Y., Zhang, Z., Wu, J., Hou, J. & Ding, G. The Exosomes Containing LINC00461 Originated from Multiple Myeloma Inhibit the Osteoblast Differentiation of Bone Mesenchymal Stem Cells via Sponging miR-324-3p. J Healthc Eng 2022, 3282860 (2022).

34. Statello, L., Guo, C.J., Chen, L.L. & Huarte, M. Gene regulation by long non-coding RNAs and its biological functions. Nature reviews. Molecular cell biology 22, 96–118 (2021).

35. Sanchez, Y. & Huarte, M. Long non-coding RNAs: challenges for diagnosis and therapies. Nucleic acid therapeutics 23, 15–20 (2013).

36. Cheetham, S.W., Gruhl, F., Mattick, J.S. & Dinger, M.E. Long noncoding RNAs and the genetics of cancer. Br J Cancer 108, 2419–2425 (2013).

37. Ling, H., Fabbri, M. & Calin, G.A. MicroRNAs and other non-coding RNAs as targets for anticancer drug development. Nature reviews. Drug discovery 12, 847–865 (2013).

38. Smith, J.S.a.D. Long Non-coding RNAs (lncRNAs) and Cancer. in Non-coding RNAs and Epigenetic Regulation of Gene Expression: Drivers of Natural Selection *| Book* (ed. Morris, K.) (Caister Academic Press, The Scripps Research Institute, La Jolla, California, USA, 2012).

39. Sangeeth, A., Malleswarapu, M., Mishra, A. & Gutti, R.K. Long Non-coding RNA Therapeutics: Recent Advances and Challenges. Curr Drug Targets 23, 1457–1464 (2022).

40. Arun, G., Diermeier, S.D. & Spector, D.L. Therapeutic Targeting of Long Non-Coding RNAs in Cancer. Trends Mol Med 24, 257–277 (2018).

41. Donlic, A., et al. Discovery of Small Molecule Ligands for MALAT1 by Tuning an RNA-Binding Scaffold. Angew Chem Int Ed Engl 57, 13242–13247 (2018).

42. Falese, J.P., Donlic, A. & Hargrove, A.E. Targeting RNA with small molecules: from fundamental principles towards the clinic. Chem Soc Rev 50, 2224–2243 (2021).

43. Liu, S.J., et al. CRISPRi-based genome-scale identification of functional long noncoding RNA loci in human cells. Science 355(2017).

44. Church, D.M., et al. Modernizing reference genome assemblies. PLoS Biol 9, e1001091 (2011).

45. Dobin, A., et al. STAR: ultrafast universal RNA-seq aligner. Bioinformatics 29, 15–21 (2013).

46. 46. Dennis, G., Jr., et al. DAVID: Database for Annotation, Visualization, and Integrated Discovery. *Genome Biol* 4, P3 (2003).

47. Zhao, W., et al. POSTAR3: an updated platform for exploring post-transcriptional regulation coordinated by RNA-binding proteins. Nucleic Acids Res 50, D287–D294 (2022).

48. Puram, S.V., et al. Cellular states are coupled to genomic and viral heterogeneity in HPV-related oropharyngeal carcinoma. Nat Genet 55, 640–650 (2023).

49. Shen, X., et al. Long Non-Coding RNA MEG3 Functions as a Competing Endogenous RNA to Regulate HOXA11 Expression by Sponging miR-181a in Multiple Myeloma. Cellular physiology and biochemistry : international journal of experimental cellular physiology, biochemistry, and pharmacology 49, 87–100 (2018).

50. Benetatos, L., et al. Promoter hypermethylation of the MEG3 (DLK1/MEG3) imprinted gene in multiple myeloma. Clin Lymphoma Myeloma 8, 171–175 (2008).

51. Guo, N., Song, Y., Zi, F., Zheng, J. & Cheng, J. Abnormal expression pattern of lncRNA H19 participates in multiple myeloma bone disease by unbalancing osteogenesis and osteolysis. Int Immunopharmacol 119, 110058 (2023).

52. Corrado, C., et al. Long Non Coding RNA H19: A New Player in Hypoxia-Induced Multiple Myeloma Cell Dissemination. International journal of molecular sciences 20(2019).

53. Pan, Y., et al. Serum level of long noncoding RNA H19 as a diagnostic biomarker of multiple myeloma.

54. *Clin Chim Acta* 480, 199-205 (2018).

54. Sun, Y., et al. Knockdown of long non-coding RNA H19 inhibits multiple myeloma cell growth via NF-kappaB pathway. Scientific reports 7, 18079 (2017).

55. Galvan-Femenia, I., et al. Multitrait genome association analysis identifies new susceptibility genes for human anthropometric variation in the GCAT cohort. J Med Genet 55, 765–778 (2018).

56. Ajith, S., et al. Position-dependent activity of CELF2 in the regulation of splicing and implications for signal-responsive regulation in T cells. RNA Biol 13, 569–581 (2016).

57. Nasiri-Aghdam, M., Garcia-Garduno, T.C. & Jave-Suarez, L.F. CELF Family Proteins in Cancer: Highlights on the RNA-Binding Protein/Noncoding RNA Regulatory Axis. International journal of molecular sciences 22(2021).

58. Yeung, Y.T., et al. CELF2 suppresses non-small cell lung carcinoma growth by inhibiting the PREX2- PTEN interaction. Carcinogenesis 41, 377–389 (2020).

59. Wang, L., et al. CELF2 is a candidate prognostic and immunotherapy biomarker in triple-negative breast cancer and lung squamous cell carcinoma: A pan-cancer analysis. J Cell Mol Med 25, 7559–7574 (2021).

60. Li, C., Mu, J., Shi, Y. & Xin, H. LncRNA CCDC26 Interacts with CELF2 Protein to Enhance Myeloid Leukemia Cell Proliferation and Invasion via the circRNA_ANKIB1/miR-195-5p/PRR11 Axis. Cell Transplant 30, 963689720986080 (2021).

61. Shi, M., et al. LncRNA-SNHG16 promotes proliferation and migration of acute myeloid leukemia cells via PTEN/PI3K/AKT axis through suppressing CELF2 protein. J Biosci 46(2021).

62. Zhao, Y., Zhou, H. & Dong, W. LncRNA RHPN1-AS1 promotes the progression of nasopharyngeal carcinoma by targeting CELF2 expression. Exp Mol Pathol 122, 104671 (2021).

63. Sarma, K., Levasseur, P., Aristarkhov, A. & Lee, J.T. Locked nucleic acids (LNAs) reveal sequence requirements and kinetics of Xist RNA localization to the X chromosome. Proc Natl Acad Sci U S A 107, 22196–22201 (2010).

64. Geary, R.S., Norris, D., Yu, R. & Bennett, C.F. Pharmacokinetics, biodistribution and cell uptake of antisense oligonucleotides. Advanced drug delivery reviews 87, 46–51 (2015).

65. Lee, J.H. & Kim, S.H. Treatment of relapsed and refractory multiple myeloma. Blood Res 55, S43–S53 (2020).

66. Radhakrishnan, S.V., et al. CD229 CAR T cells eliminate multiple myeloma and tumor propagating cells without fratricide. Nature communications 11, 798 (2020).

67. Vander Mause, E.R., et al. Systematic single amino acid affinity tuning of CD229 CAR T cells retains efficacy against multiple myeloma and eliminates on-target off-tumor toxicity. Sci Transl Med 15, eadd7900 (2023).

68. O’Neal, J., et al. CS1 CAR-T targeting the distal domain of CS1 (SLAMF7) shows efficacy in high tumor burden myeloma model despite fratricide of CD8+CS1 expressing CAR-T cells. Leukemia 36, 1625–1634 (2022).

69. Cooper, M.L. & DiPersio, J.F. Chimeric antigen receptor T cells (CAR-T) for the treatment of T-cell malignancies. Best Pract Res Clin Haematol 32, 101097 (2019).

70. Zhou, M., et al. Identification and validation of potential prognostic lncRNA biomarkers for predicting survival in patients with multiple myeloma. J Exp Clin Cancer Res 34, 102 (2015).

71. Dong, H., et al. Upregulation of lncRNA NR_046683 Serves as a Prognostic Biomarker and Potential Drug Target for Multiple Myeloma. Frontiers in pharmacology 10, 45 (2019).

72. Ronchetti, D., et al. A compendium of long non-coding RNAs transcriptional fingerprint in multiple myeloma. Scientific reports 8, 6557 (2018).

73. Zhang, L., Liu, X., Zhang, X. & Chen, R. Identification of important long non-coding RNAs and highly recurrent aberrant alternative splicing events in hepatocellular carcinoma through integrative analysis of multiple RNA-Seq datasets. Mol Genet Genomics 291, 1035–1051 (2016).

74. Yang, Y., Cheng, Y., Mou, Y., Tang, X. & Mu, X. Natural Antisense Long Noncoding RNA HHIP-AS1 Suppresses Non-Small-Cell Lung Cancer Progression by Increasing HHIP Stability via Interaction with CELF2. Crit Rev Eukaryot Gene Expr 33, 67–77 (2022).

75. Xie, S.C., et al. LncRNA CRNDE facilitates epigenetic suppression of CELF2 and LATS2 to promote proliferation, migration and chemoresistance in hepatocellular carcinoma. Cell death & disease 11, 676 (2020).

76. Wang, D., et al. GAS5 knockdown alleviates spinal cord injury by reducing VAV1 expression via RNA binding protein CELF2. Scientific reports 11, 3628 (2021).

77. Vejdandoust, F., Moosavi, R., Fattahi Dolatabadi, N., Zamani, A. & Tabatabaeian, H. MIMT1 and LINC01550 are uncharted lncRNAs down-regulated in colorectal cancer. Int J Exp Pathol 104, 107–116 (2023).

78. Tuo, H., Liu, R., Wang, Y., Yang, W. & Liu, Q. Hypoxia-induced lncRNA MRVI1-AS1 accelerates hepatocellular carcinoma progression by recruiting RNA-binding protein CELF2 to stabilize SKA1 mRNA. World journal of surgical oncology 21, 111 (2023).

79. Tian, Y., Sun, L. & Qi, T. Long noncoding RNA GAS5 ameliorates chronic constriction injury induced neuropathic pain in rats by modulation of the miR-452-5p/CELF2 axis. Can J Physiol Pharmacol 98, 870–877 (2020).

80. Guo, D., Zhang, A., Suo, M., Wang, P. & Liang, Y. ELK1-Induced upregulation of long non-coding TNK2- AS1 promotes the progression of acute myeloid leukemia by EZH2-mediated epigenetic silencing of CELF2. Cell Cycle 22, 117–130 (2023).

81. Lai, S., et al. N6-methyladenosine-mediated CELF2 regulates CD44 alternative splicing affecting tumorigenesis via ERAD pathway in pancreatic cancer. Cell Biosci 12, 125 (2022).

82. Chatrikhi, R., et al. RNA Binding Protein CELF2 Regulates Signal-Induced Alternative Polyadenylation by Competing with Enhancers of the Polyadenylation Machinery. Cell reports 28, 2795–2806 e2793 (2019).

83. Mallory, M.J., et al. Reciprocal regulation of hnRNP C and CELF2 through translation and transcription tunes splicing activity in T cells. Nucleic Acids Res 48, 5710–5719 (2020).

84. Subramaniam, D., et al. Translation inhibition during cell cycle arrest and apoptosis: Mcl-1 is a novel target for RNA binding protein CUGBP2. Am J Physiol Gastrointest Liver Physiol 294, G1025–1032 (2008).

85. Li, J.H., et al. Discovery of Protein-lncRNA Interactions by Integrating Large-Scale CLIP-Seq and RNA-Seq Datasets. Front Bioeng Biotechnol 2, 88 (2014).

86. Davidovich, C., Zheng, L., Goodrich, K.J. & Cech, T.R. Promiscuous RNA binding by Polycomb repressive complex 2. Nature structural & molecular biology 20, 1250–1257 (2013).

87. Jiang, C., et al. Identifying and functionally characterizing tissue-specific and ubiquitously expressed human lncRNAs. Oncotarget 7, 7120–7133 (2016).

88. Chen, X. & Sun, Z. Novel lincRNA Discovery and Tissue-Specific Gene Expression across 30 Normal Human Tissues. Genes (Basel) 12(2021).

89. Dowerah, D., et al. Design of LNA Analogues Using a Combined Density Functional Theory and Molecular Dynamics Approach for RNA Therapeutics. ACS Omega 8, 22382–22405 (2023).

90. Lundin, K.E., et al. Biological activity and biotechnological aspects of locked nucleic acids. Adv Genet 82, 47–107 (2013).

91. Kuespert, S., et al. Antisense Oligonucleotide in LNA-Gapmer Design Targeting TGFBR2-A Key Single Gene Target for Safe and Effective Inhibition of TGFbeta Signaling. International journal of molecular sciences 21(2020).

92. Dhuri, K., et al. Antisense Oligonucleotides: An Emerging Area in Drug Discovery and Development. J Clin Med 9(2020).

94. 93. (2024), U.S.N.L.o.M. Clinicaltrials.gov. (2024).

94. Maruyama, R. & Yokota, T. Knocking Down Long Noncoding RNAs Using Antisense Oligonucleotide Gapmers. Methods Mol Biol 2176, 49–56 (2020).

95. Zhou, T., Kim, Y. & MacLeod, A.R. Targeting Long Noncoding RNA with Antisense Oligonucleotide Technology as Cancer Therapeutics. Methods Mol Biol 1402, 199–213 (2016).

96. Goyal, B., et al. Diagnostic, prognostic, and therapeutic significance of long non-coding RNA MALAT1 in cancer. Biochim Biophys Acta Rev Cancer 1875, 188502 (2021).

97. Taiana, E., et al. Long non-coding RNA NEAT1 targeting impairs the DNA repair machinery and triggers anti-tumor activity in multiple myeloma. Leukemia 34, 234–244 (2020).

